# Structure-Function Relationship of the Ryanodine Receptor Cluster Network in Sinoatrial Node Cells

**DOI:** 10.1101/2024.10.09.617454

**Authors:** Alexander V Maltsev, Valeria Ventura Subirachs, Oliver Monfredi, Magdalena Juhaszova, Pooja Ajay Warrier, Shardul Rakshit, Syevda Tagirova, Anna V Maltsev, Michael D Stern, Edward G Lakatta, Victor A Maltsev

**Affiliations:** National Institute on Aging, NIH, Baltimore, MD 21224, USA; Department of Cardiovascular Electrophysiology, The Johns Hopkins Hospital, Baltimore, MD, 21287, USA; School of Mathematics, Queen Mary University of London, London, UK

**Keywords:** β adrenergic stimulation, numerical model, pacemaker function, ryanodine receptor, sinoatrial node

## Abstract

The rate of spontaneous action potentials (APs) generated by sinoatrial node cells (SANC) is regulated by local Ca^2+^ release (LCR) from the sarcoplasmic reticulum via Ca^2+^ release channels (ryanodine receptors, RyRs). LCR events propagate and self-organize within the network of RyR clusters (Ca release units, CRUs) via Ca-induced-Ca-release (CICR) that depends on CRU sizes and locations: while larger CRUs generate stronger release signals, the network’s topology governs signal diffusion and propagation. This study used super-resolution structured illumination microscopy to image the 3D network of CRUs in rabbit SANC. The peripheral CRUs formed a spatial mesh, reflecting the cell surface geometry. Two distinct subpopulations of CRUs were identified within each cell, with size distributions conforming to a two component Gamma mixture model. Furthermore, neighboring CRUs exhibited repulsive behavior. Functional properties of the CRU network were further examined in a novel numerical SANC model developed using our experimental data. Model simulations revealed that heterogeneities in both CRU sizes and locations *facilitate* CICR and increase AP firing rate in a cooperative manner. However, these heterogeneities reduce the effect of β-adrenergic stimulation in terms of its *relative change* in AP firing rate. The presence of heterogeneities in both sizes and locations allows SANC to reach higher *absolute AP firing rates* during β-adrenergic stimulation. Thus, the CICR facilitation by heterogeneities in CRU sizes and locations regulates and optimizes cardiac pacemaker cell operation under various physiological conditions. Dysfunction of this optimization could be a key factor in heart rate reserve decline in aging and disease.

## 1. Introduction

Sinoatrial node cells (SANC) located in the posterosuperior right atrium are specialized to generate rhythmic electrical impulses that drive cardiac contractions [1,2]. SANC dysfunction can lead to sick sinus syndrome associated with a variety of life-threatening arrhythmias [3-6]. Generation of normal rhythmic impulses by SANC is executed via coupled signaling of both cell membrane ion channels and Ca^2+^ cycling, known as the coupled-clock system [7]. A vital component of this system is the sarcoplasmic reticulum (SR), the main intracellular Ca^2+^ store consisting of two parts: (i) the network SR that uptakes cytosolic Ca^2+^ via a Ca^2+^ pump and (ii) the junctional SR (JSR) that embeds clusters of 10-200 Ca^2+^ release channels (ryanodine receptors, RyRs [8]), forming functional couplons or Ca^2+^ release units (CRUs) linked to L-type channels [9]. While RyRs release Ca^2+^ from JSR, their openings are also activated by Ca, creating a positive (i.e. explosive) feedback, known as Ca-induced-Ca release (CICR) [10]. Thus, RyRs within a CRU tend to release Ca^2+^ in synchrony, creating an elementary Ca^2+^ signal, known as Ca^2+^ spark [11,12]. In SANC under normal conditions, a Ca^2+^ spark, in turn, can interact with neighboring CRUs via CICR creating a series of locally propagating sparks, observed as Ca^2+^ wavelets, known as local Ca^2+^ release (LCR) (review [7]). The LCRs, in turn, activate electrogenic Na/Ca exchanger embedded in cell surface membrane that accelerates diastolic depolarization [7,13-15]. The diastolic depolarization is further accelerated by a positive feedback mechanism that includes voltage-activated L-type Ca^2+^ channels which not only depolarize the membrane, but also bring more Ca^2+^ into the cell. Thus, the Ca^2+^ influx brings more “fuel” to further explode CICR and diastolic depolarization, ensuring robust action potential (AP) ignition [16].

In this complex chain of signaling events CICR plays a critical role, and its efficiency must strongly depend on the spatial organization of CRU network, i.e. the distributions of CRU sizes and locations. What is known thus far about the CRU network in SANC and what is still missing? RyRs are indeed clustered under the cell membrane as observed in immunofluorescence images of SANC of various species [17,18]. Early simple numerical models of CRU networks in SANC considered identical CRUs in a perfect square grid [19,20]. These models demonstrated an important role of CICR in cardiac pacemaking, e.g. the emergence of a Ca^2+^ clock and robust AP firing (via stabilizing diastolic Na/Ca exchanger current [20]). Subsequentially distributions of CRU sizes and nearest neighbor distances (NND) were measured in 2D in tangential sections of confocal microscopy images and included in a 3D model of SANC [21]. That study raised important questions of CRU function such as “when can a Ca^2+^ spark jump?”, i.e. when CICR propagate among neighboring CRUs. CICR and AP firing emerged in the model only when larger sized CRUs were mixed with another population of smaller CRUs that provided bridges for CICR propagation, indicating importance of CRU size heterogeneity. More recent theoretical studies have demonstrated that disorder in CRU locations confers robustness but reduces flexibility in heart pacemaking [22], showing importance of heterogeneity in CRU distances. This was shown, however, in a SANC model with CRUs of identical sizes and hypothetical CRU repulsion, whereas precise organization of the CRU network in 3D and the functional role of heterogeneity of CRU sizes remain unknown. Thus, it is still not understood how the Ca^2+^ clock and the coupled clock system emerge from the scale and organization of RyRs and CRUs towards the whole cell function [23,24].

The present study approaches this problem by combining experimental methods and numerical modelling. We imaged the precise organization of the CRU network (both sizes and locations) in 3D in rabbit SANC by super-resolution structured illumination microscopy (SIM), that provided larger optical (about twice) resolution compared to confocal imaging [25]. Our results revealed that CRU sizes follow a mixture of two Gamma distributions, suggesting distinct subpopulations of smaller and larger CRUs. Furthermore, NNDs of RyR clusters exhibit a power law distribution at short distances, indicating repulsive interactions among CRUs that would restrain CICR and Ca^2+^ signal propagation. Based on these findings, we developed a new mathematical SANC model featuring CRUs of heterogeneous sizes and locations and performed model simulations with different CRU networks to obtain deeper insights. The simulations demonstrated that complex and flexible structure of the RyR network, including partial disorder, represents a new pacemaker regulatory mechanism that optimizes cardiac pacemaker cell operation under various physiological conditions.

## 2. Materials and Methods

### 2.1. Enzymatic isolation of individual SANC

SANC were isolated from male rabbits in accordance with NIH guidelines for the care and use of animals, protocol # 457-LCS-2024 as previously described [26]. New Zealand White rabbits (Charles River Laboratories, USA) weighing 2.8–3.2 Kg were anesthetized with sodium pentobarbital (50–90 mg/kg). The heart was removed quickly and placed in solution containing (in mM): 130 NaCl, 24 NaHCO_3_, 1.2 NaH_2_PO_4_, 1.0 MgCl_2_, 1.8 CaCl_2_, 4.0 KCl, 5.6 glucose equilibrated with 95% O_2_ / 5% CO_2_ (pH 7.4 at 35°C). The SAN region was cut into small strips (∼1.0 mm wide) perpendicular to the crista terminalis and excised. The final SA node preparation, which consisted of SA node strips attached to the small portion of crista terminalis, was washed twice in nominally Ca^2+^-free solution containing (in mM): 140 NaCl, 5.4 KCl, 0.5 MgCl2, 0.33 NaH_2_PO_4_, 5 HEPES, 5.5 glucose, (pH=6.9) and incubated on shaker at 35°C for 30 min in the same solution with the addition of elastase type IV (0.6 mg/ml; Sigma, Chemical Co.), collagenase type 2 (0.8 mg/ml; Worthington, NJ, USA), Protease XIV (0.12 mg/ml; Sigma, Chemical Co.), and 0.1% bovine serum albumin (Sigma, Chemical Co.). The SA node preparation was next placed in modified “Kraftbruhe” solution, containing (in mM): 70 potassium glutamate, 30 KCl, 10 KH_2_PO_4_, 1 MgCl_2_, 20 taurine, 10 glucose, 0.3 EGTA, and 10 HEPES (titrated to pH 7.4 with KOH), and kept at 4°C for 1h in KB solution containing 50 mg/ml polyvinylpyrrolidone (PVP, Sigma, Chemical Co.). Finally, cells were dispersed from the SA node preparation by gentle pipetting in the “Kraftbruhe” solution and stored at 4°C.

### 2.2. Immunolabeling of RyRs

Isolated cells were fixed with 2% formaldehyde in phosphate buffered saline (PBS, Sigma) and then permeabilized with 0.5-1% Triton X-100/PBS. Nonspecific cross-reactivity was blocked by incubating the samples for 4 h in 1% BSA/PBS (Jackson ImmunoResearch, West Grove, PA, USA). The cells were then incubated with primary antibodies anti-RyR (IgG1, clone C3-33, Afinity BioReagents, Golden, CO, or AP20325PU-N (Origene), for details see our database at https://doi.org/10.7910/DVN/N0OLGI). The antibodies were diluted in 1%BSA/PBS overnight, washed in PBS, and then incubated with appropriate fluorescence label-conjugated secondary antibodies (Jackson ImmunoResearch, West Grove, PA).

### 2.3. Structured illumination microscopy (SIM)

SIM uses a sharply patterned light source to improve spatial resolution in fluorescence microscopy [27]. SIM can be used to study the structure and function of cells in great detail, as it can provide images with a resolution that is higher than that of traditional microscopy techniques. It is particularly useful for studying cells and tissues that have fine structural features, such as the SANC, as we report here. Experiments with RyR detection in fixed cells were performed in the experimental instant SIM setup in the Hari Shroff’s laboratory at the National Institute of Biomedical Imaging and Bioengineering, as described elsewhere [27-29]. The instant SIM is an implementation of SIM well-suited for high-speed imaging, as the images are processed optically rather than computationally to improve resolution ∼1.4x. SANC were imaged via an oil x100, 1.49 NA objective and 488 nm excitation; in-house software controlled optimal laser power and positioning (sectioning) of cell preparation. The system allowed imaging at resolutions as low as 145 nm laterally and 320 nm axially. 3D sectioning was performed with a pixel size of 55.5 × 55.5 nm in *xy* and *z* stacking of 150 nm.

### 2.4. Segmentation of peripheral RyR clusters in 3D

Our dataset consisted of 31 rabbit SANC which included an aggregated total of 70,982 RyR clusters. We present a novel algorithm for high-throughput segmentation of peripheral ryanodine receptor (RyR) clusters in 3D SANC (Figure 1). The preprocessing procedure included the following steps in order: a 3D median filter to remove salt-and-pepper noise, signal normalization to standardize the intensity range, and finally, 3D contrast-limited adaptive histogram equalization (CLAHE) [30] to enhance local contrast while preserving overall image structure and global contrast. The segmentation was then performed using the 3D StarDist neural network [31], which directly predicts star-convex polyhedra representations for each RyR cluster without relying on the watershed algorithm. 3D StarDist learns to separate partially overlapping or contacting RyR clusters by predicting appropriate polyhedra shapes based on the trained model. The ground truth data used for training 3D StarDist was initially generated using Squassh [32], an open-source software algorithm in ImageJ Fiji, which is part of the MOSAIC suite that utilizes globally optimal detection and segmentation methodologies [33] and incorporates corrections for the microscope’s point spread function [34,35]. The Squassh segmentation results were then fine-tuned using the Adaptive Watershed tool in ORS Dragonfly [36] and further refined by manual editing in ORS Dragonfly to ensure accurate separation of RyR clusters (Figure 2). The final trained model for 3D StarDist achieved an Intersection over Union (IoU) score of 0.68.

**Figure 1.**
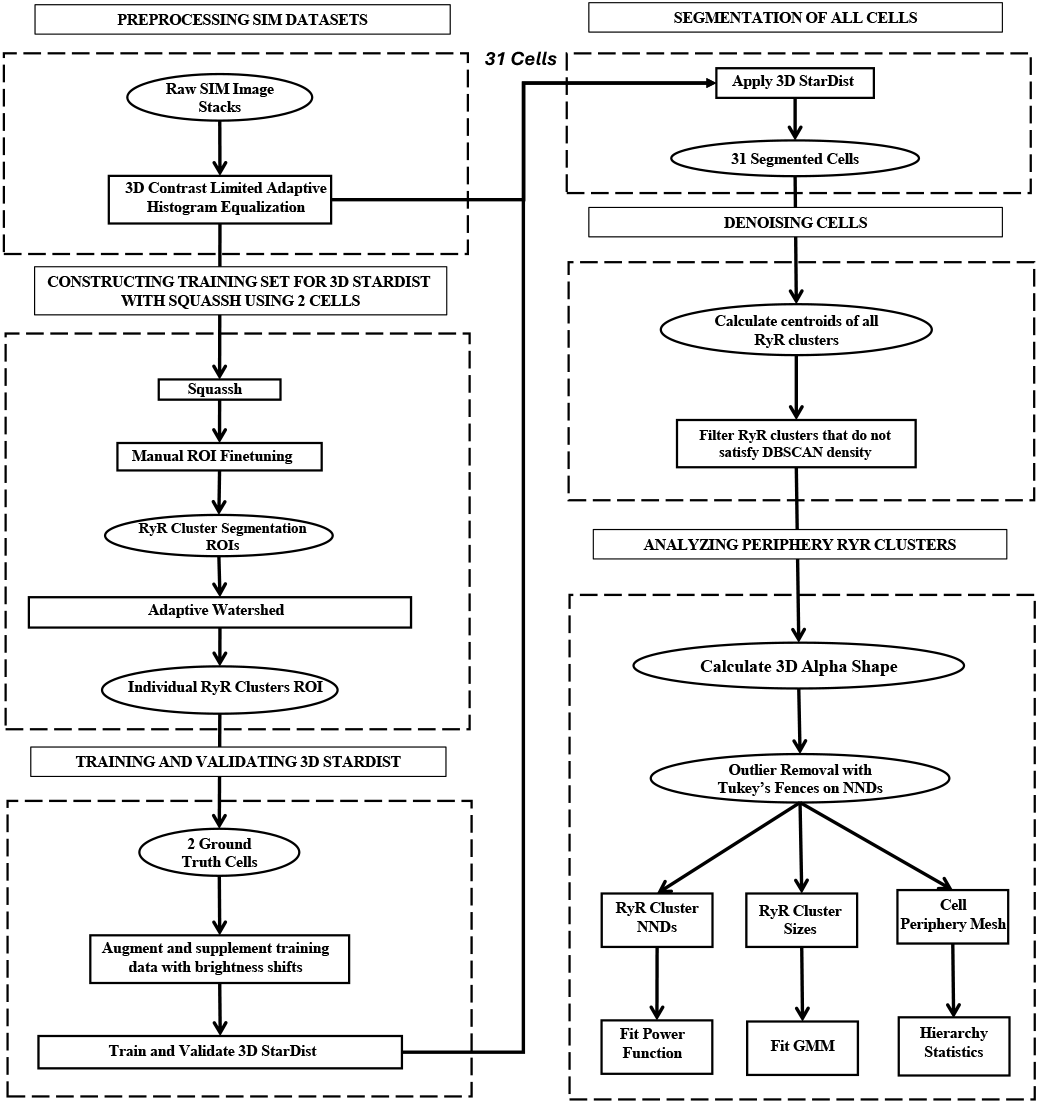
Flowchart of the peripheral RyR cluster analysis algorithm. Presented is our image processing algorithm for the precise segmentation and analysis of peripheral RyR clusters in 3D SIM imaging data in SANC. Initially, the data undergoes cropping and contrast enhancement through 3D CLAHE. The main segmentation process is conducted using the 3D StarDist neural network, which is trained using ground truth data generated through the Squassh software and refined via adaptive watershed. Following segmentation, RyR clusters that are part of the cell are identified and extracted using the density-based spatial clustering of applications with noise (DBSCAN) algorithm, which detects high-density clusters within the dataset. The culmination of this process is the generation of a 3D alpha shape, encapsulating the spatial distribution of the RyR clusters on the periphery of SANC. The peripheral RyR cluster sizes, distances, and their alpha shape mesh is exported for further statistical processing.

**Figure 2.**
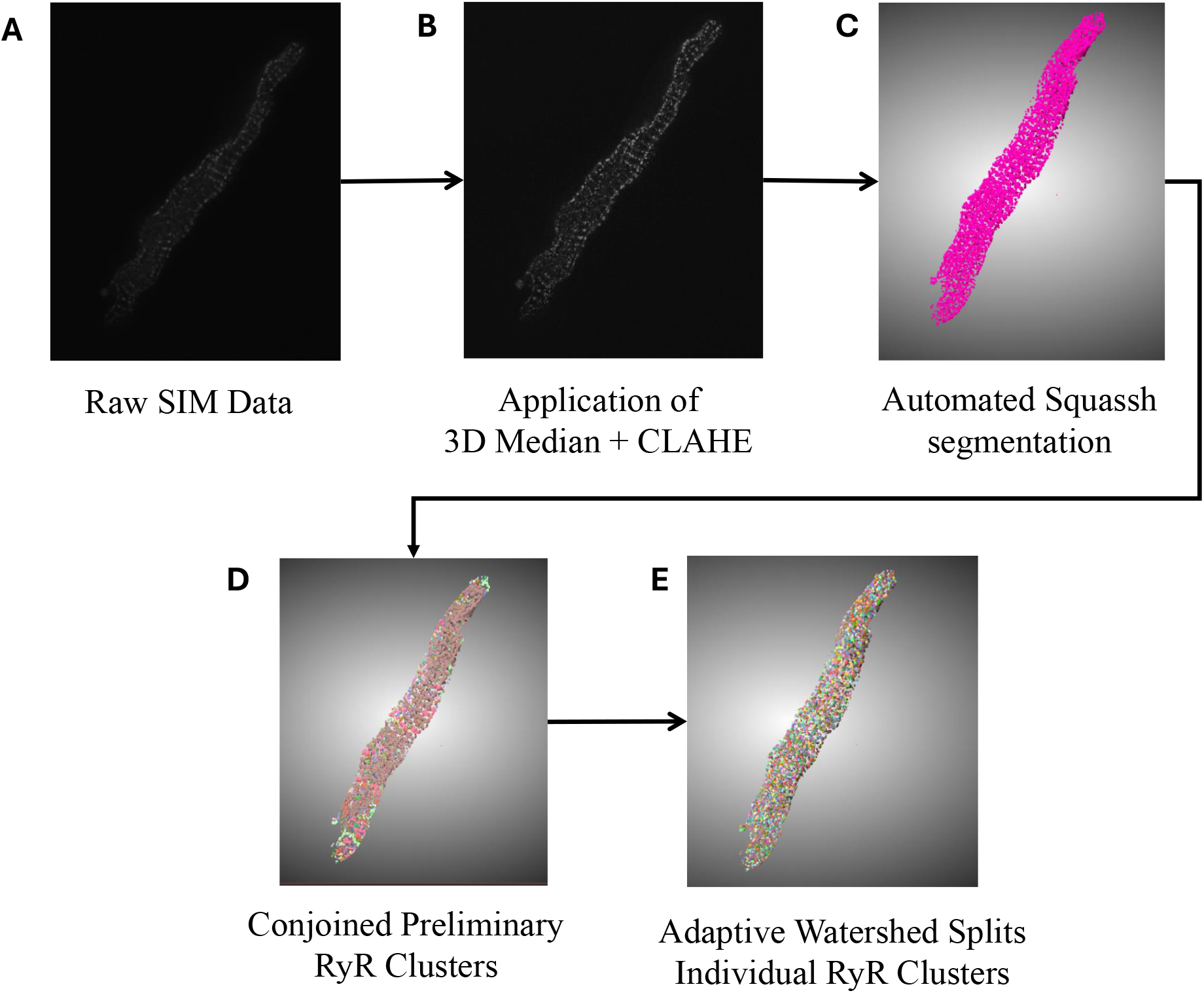
Visual representation of transformations and segmentation of flowchart. (**A**): the initial, unprocessed SIM image data (in 3 dimensions) of RyR immunofluorescence in SANC, serving as the primary source for image analysis. (**B**): the enhanced and denoised raw SIM data. (**C**): the preliminary segmentation from Squassh with a purple ROI. (**D**): the stage where distinct RyR clusters are isolated, establishing boundaries between adjoining regions using a 3D 26-connectivity strategy. (**E**): the Watershed algorithm’s role in further refining the segmentation, capable of isolating individual RyR clusters even in complex spatial arrangements.

To further refine the segmentation of RyR clusters and remove false positives after 3D StarDist segmentation, DBSCAN (Density-Based Spatial Clustering of Applications with Noise) clustering was used on the centroid data of the segmented RyR clusters. DBSCAN is a density-based clustering algorithm that groups together points that are closely packed, marking points that are in low-density regions as outliers. The key parameters for DBSCAN were ∈ and *min_samples*. ϵ determines the maximum distance between two points for them to be considered as part of the same cluster or in our case, the imaged SANC. ϵ was set using the median absolute deviation (MAD) method with a scaling factor *c*.

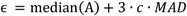

By using the scaled MAD and the median, the ϵ remained robust against the influence of extreme values:

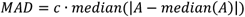

where A represents the set of nearest neighbor distances. Furthermore, the scaling factor c was defined as:

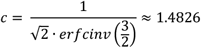

where *erfcinv* is the inverse complementary error function. This adaptation ensures that the scaled MAD approximates the standard deviation when the data follows a normal distribution. The second parameter *min_samples* is the minimum number of points required to form a dense region and it was set to 3. If a point has at least *min_samples* within its ϵ-neighborhood (including itself), it is considered a core point. Points that are not core points but are within the ϵ-neighborhood of a core point are called border points. Points that are neither core points nor border points are classified as noise or outliers.

The largest cluster deduced by DBSCAN (which is the cell itself) was further refined by computing its alpha shape, which captured the peripheral RyR clusters. The optimal alpha value was determined by iterating through a range of values and selecting the one that maximizes the density of points on the surface of the alpha shape. Finally, the algorithm generated various data and visualizations, including the alpha shape, surface area, volume, centroid data, volume data, voxel lists, cluster labels, adjacency matrix, and nearest neighbor distances, for further analysis. An additional outlier test was performed by taking Tukey’s Fences where extreme nearest neighbor values are filtered out by keeping only the interval:

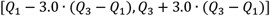

The visualizations for all 31 cells are presented in Figure 3 and mouse-interactive HTML files constructed with *plotly* Python library are published in the GitHub https://github.com/alexmaltsev/SANC/tree/main/3D%20Visualizations. Each cell can be individually opened and rotated (with zoom-in or -out) by the reader. We also published in the GitHub our code to use 3D StarDist and all code for data application and processing. See Code Availability section for more details.

**Figure 3.**
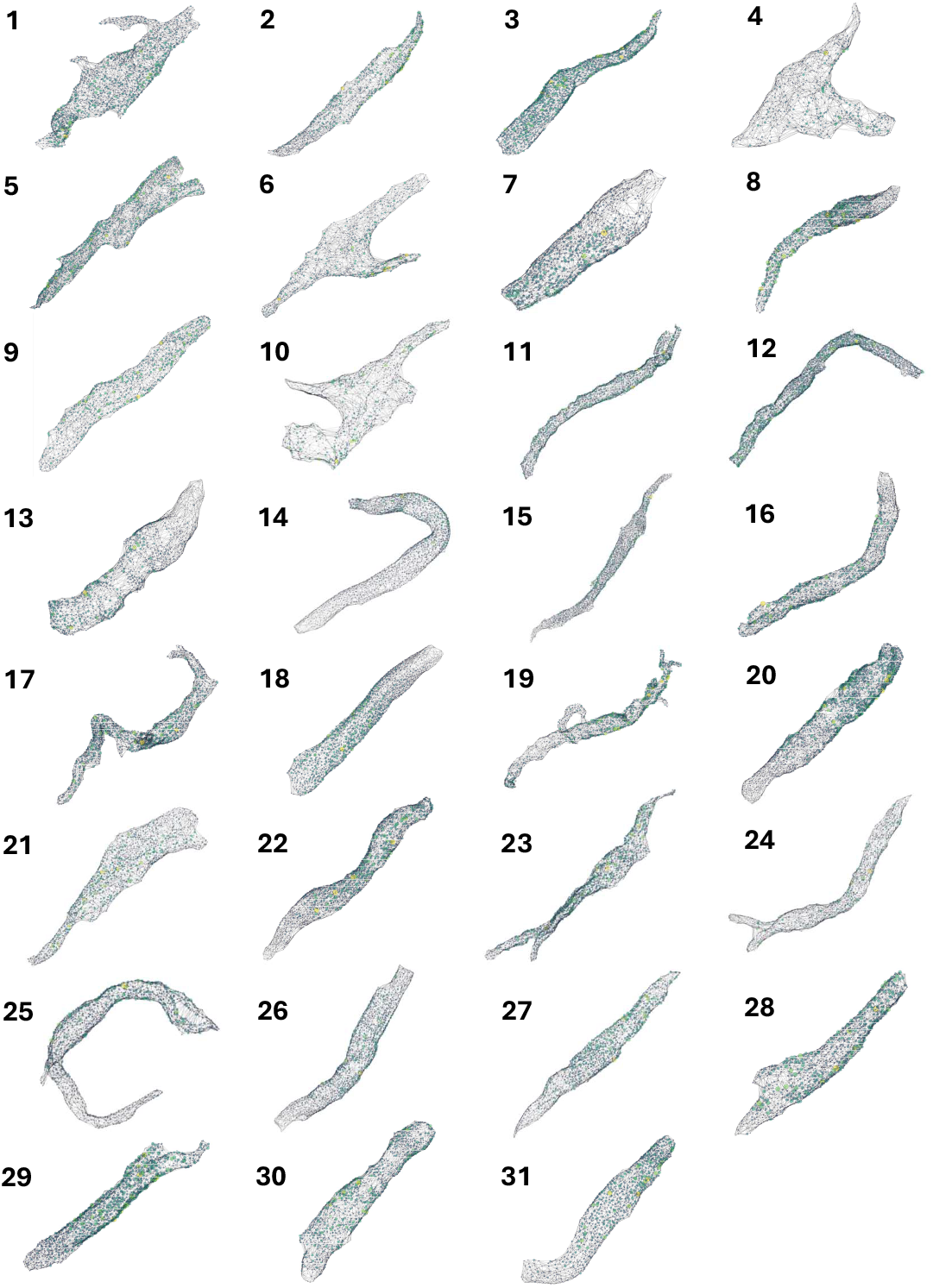
3D visualization of RyR cluster distribution in all 31 rabbit SANC analyzed in the study. Each cell is represented by a unique number (1-31) corresponding to the cell identifiers used in Tables 1 and 2. The visualizations depict the spatial distribution of RyR clusters within each cell, with clusters represented as colored points. The color intensity indicates the relative size of the clusters, with brighter colors (yellow to green) representing larger clusters and dimmer colors (blue to gray) representing smaller clusters. These visualizations provide a comprehensive overview of the RyR cluster organization across the entire sample set, allowing for visual comparison of cluster distributions and densities among cells of different morphologies. Cells are shown at various zoom levels to closely fit the panel size. 3D representation of each cell with its scale bar is available in html format in the GitHub https://github.com/alexmaltsev/SANC/tree/main/3D%20Visualizations. Each html file can be opened in a web browser for computer mouse-interactive view (rotation and zoom-in and -out).

## 3. Results

### 3.1. Two subpopulations of RyR cluster sizes

The distributions of RyR cluster sizes in individual SANC were analyzed, with results shown in Table 1. On average we detected 2285 clusters per cell and overall mean of RyR cluster sizes was 0.18 μm^3^ with standard deviation 0.13 μm^3^. The distributions were plotted as histograms with 50 bins and the existence of two right side skewed subpopulations was visually apparent with four representative cells shown in Figure 4. Thus, a Gamma Mixture Model (GMM) was used to model this distribution. A Gamma distribution is often used to model distributions that are asymmetric, with a longer tail on the right side. A GMM is a probabilistic model that assumes the data is generated from a mixture of several Gamma distributions, in our case two distributions. Each component in the mixture is a Gamma distribution with its own parameters, and the mixture is weighted by coefficients that sum to 1. The PDF for a GMM with two components in our case is:

**Table 1.** Morphological and RyR cluster characteristics of rabbit sinoatrial node cells. Detailed statistics for 31 rabbit sinoatrial node cells, including their surface area, volume, number of RyR clusters, and metrics related to RyR cluster distances and sizes. The data encompasses a wide range of cellular morphologies and RyR cluster distributions. Notably, cells 3, 8, 14, 24, and 27 (in Figure 3) are included in this comprehensive dataset, allowing for direct comparison of their characteristics with the broader cell population.

| Cell | Cell Statistics |  |  | RyR Cluster Distances |  | RyR Cluster Sizes |  |
| --- | --- | --- | --- | --- | --- | --- | --- |
| | Surface Area ($\mu\text{m}^2$) | Volume ($\mu\text{m}^3$) | #RYR Clusters | Mean (nm) | SD (nm) | Mean ($\mu\text{m}^3$) | SD ($\mu\text{m}^3$) |
| 1 | 2494 | 2886 | 2775 | 586.78 | 191.25 | 0.21 | 0.17 |
| 2 | 2166 | 3050 | 1871 | 633.27 | 210.61 | 0.16 | 0.12 |
| 3 | 2698 | 5234 | 2035 | 722.19 | 218.38 | 0.26 | 0.15 |
| 4 | 2177 | 3695 | 1359 | 657.48 | 239.13 | 0.13 | 0.09 |
| 5 | 3182 | 4777 | 3341 | 579.93 | 212.91 | 0.17 | 0.17 |
| 6 | 2708 | 3868 | 1901 | 647.02 | 233.15 | 0.12 | 0.08 |
| 7 | 2435 | 6409 | 1566 | 695.58 | 243.85 | 0.19 | 0.14 |
| 8 | 1923 | 3119 | 2085 | 627.35 | 185.29 | 0.19 | 0.15 |
| 9 | 1583 | 1992 | 1573 | 617.39 | 209.54 | 0.19 | 0.16 |
| 10 | 2879 | 5088 | 1675 | 705.36 | 268.41 | 0.12 | 0.09 |
| 11 | 2300 | 2722 | 3046 | 555.98 | 174.95 | 0.20 | 0.18 |
| 12 | 3175 | 4822 | 3603 | 600.26 | 179.40 | 0.20 | 0.13 |
| 13 | 3097 | 8328 | 1976 | 702.53 | 229.66 | 0.14 | 0.09 |
| 14 | 2841 | 4579 | 2648 | 669.51 | 196.30 | 0.17 | 0.11 |
| 15 | 2129 | 2150 | 3243 | 530.22 | 164.79 | 0.16 | 0.13 |
| 16 | 2240 | 3431 | 2470 | 587.66 | 204.89 | 0.18 | 0.16 |
| 17 | 2139 | 2238 | 2577 | 570.93 | 185.03 | 0.19 | 0.16 |
| 18 | 2457 | 4984 | 2074 | 704.40 | 207.67 | 0.18 | 0.12 |
| 19 | 2688 | 2813 | 2575 | 606.69 | 196.25 | 0.17 | 0.15 |
| 20 | 2457 | 3735 | 2201 | 648.78 | 203.09 | 0.19 | 0.15 |
| 21 | 5256 | 15899 | 3364 | 736.34 | 253.06 | 0.20 | 0.14 |
| 22 | 2195 | 3898 | 2548 | 557.56 | 181.65 | 0.16 | 0.13 |
| 23 | 2318 | 2772 | 2292 | 633.41 | 194.55 | 0.23 | 0.18 |
| 24 | 2474 | 3655 | 2328 | 605.15 | 212.17 | 0.12 | 0.09 |
| 25 | 1983 | 2200 | 2205 | 596.14 | 191.90 | 0.18 | 0.14 |
| 26 | 2908 | 6224 | 2370 | 692.24 | 210.53 | 0.18 | 0.12 |
| 27 | 1877 | 3121 | 1639 | 676.19 | 209.62 | 0.18 | 0.14 |
| 28 | 1567 | 2278 | 1633 | 596.97 | 195.09 | 0.17 | 0.13 |
| 29 | 2564 | 4668 | 1987 | 694.89 | 212.81 | 0.19 | 0.12 |
| 30 | 3117 | 8181 | 2406 | 671.59 | 238.41 | 0.20 | 0.15 |
| 31 | 1640 | 2553 | 1483 | 687.51 | 198.09 | 0.22 | 0.14 |
| AVERAGE | 2505 | 4367 | 2285 | 638.62 | 208.14 | 0.18 | 0.13 |

**Table 2.** RyR cluster distribution analysis in rabbit SANC. Detailed analytical results for the 31 rabbit SANC, focusing on key aspects of RyR cluster distribution: repulsion (b) and Gamma Mixture Model (GMM) parameters (α1, β1, α2, β2, *π*). The repulsion parameter b indicates the strength of spatial repulsion between RyR clusters. The GMM parameters describe the distribution of RyR cluster sizes. Notably, cells 3, 8, 14, 24, and 27 in previous figures are included in this dataset, allowing for a more comprehensive understanding of their RyR cluster characteristics in the context of the entire cell population. The aggregated results refer to the results that are calculated from all RyR clusters in all 31 cells in a single plot.

|  | Repulsion | Gamma Mixture Model (GMM) |  |  |  |  |
| --- | --- | --- | --- | --- | --- | --- |
| Cell | $b$ | $\alpha_1$ | $\beta_1$ | $\alpha_2$ | $\beta_2$ | $\pi$ |
| 1 | 2.59 | 4.20 | 142.86 | 2.43 | 9.52 | 0.19 |
| 2 | 5.87 | 4.05 | 142.86 | 2.28 | 12.99 | 0.12 |
| 3 | 3.82 | 2.48 | 43.48 | 5.39 | 17.86 | 0.16 |
| 4 | 2.51 | 4.79 | 111.11 | 2.42 | 16.67 | 0.17 |
| 5 | 7.76 | 4.51 | 142.86 | 2.12 | 8.55 | 0.34 |
| 6 | 4.65 | 5.20 | 125.00 | 2.77 | 20.83 | 0.13 |
| 7 | 3.01 | 3.47 | 100.00 | 3.04 | 13.51 | 0.20 |
| 8 | 2.34 | 4.30 | 111.11 | 2.23 | 10.10 | 0.19 |
| 9 | 4.05 | 3.93 | 125.00 | 2.19 | 9.17 | 0.23 |
| 10 | 3.13 | 5.05 | 111.11 | 2.62 | 18.87 | 0.18 |
| 11 | 3.79 | 4.57 | 142.86 | 1.99 | 7.87 | 0.22 |
| 12 | 5.76 | 2.03 | 35.71 | 3.36 | 14.29 | 0.23 |
| 13 | 4.2 | 4.45 | 125.00 | 2.99 | 19.61 | 0.10 |
| 14 | 4.07 | 1.94 | 12.99 | 8.03 | 35.71 | 0.74 |
| 15 | 5.71 | 4.38 | 111.11 | 2.18 | 11.90 | 0.14 |
| 16 | 5.12 | 4.67 | 142.86 | 1.80 | 8.13 | 0.20 |
| 17 | 3.21 | 4.18 | 125.00 | 2.14 | 9.17 | 0.22 |
| 18 | 3.9 | 1.65 | 10.53 | 8.60 | 34.48 | 0.69 |
| 19 | 3.81 | 4.15 | 111.11 | 2.10 | 9.90 | 0.21 |
| 20 | 3.2 | 4.55 | 142.86 | 2.24 | 10.42 | 0.12 |
| 21 | 4.1 | 4.21 | 111.11 | 2.99 | 12.66 | 0.18 |
| 22 | 5.35 | 3.64 | 100.00 | 2.21 | 11.63 | 0.22 |
| 23 | 3.96 | 3.86 | 90.91 | 2.60 | 9.43 | 0.18 |
| 24 | 5.5 | 2.57 | 33.33 | 4.12 | 22.22 | 0.61 |
| 25 | 3.21 | 8.62 | 333.33 | 1.91 | 9.71 | 0.12 |
| 26 | 5.74 | 1.83 | 11.90 | 7.22 | 29.41 | 0.72 |
| 27 | 3.23 | 4.37 | 125.00 | 2.74 | 12.20 | 0.21 |
| 28 | 4.45 | 2.69 | 55.56 | 2.79 | 12.66 | 0.30 |
| 29 | 4.59 | 1.89 | 11.24 | 9.14 | 37.04 | 0.69 |
| 30 | 4.19 | 3.96 | 125.00 | 2.89 | 11.76 | 0.22 |
| 31 | 3.03 | 1.52 | 8.26 | 8.11 | 29.41 | 0.62 |
| AVERAGE | 4.19 | 3.80 | 100.68 | 3.54 | 16.05 | 0.29 |
| AGGREGATED | 5.15 | 4.35 | 125.00 | 2.22 | 10.64 | 0.16 |

**Figure 4.**
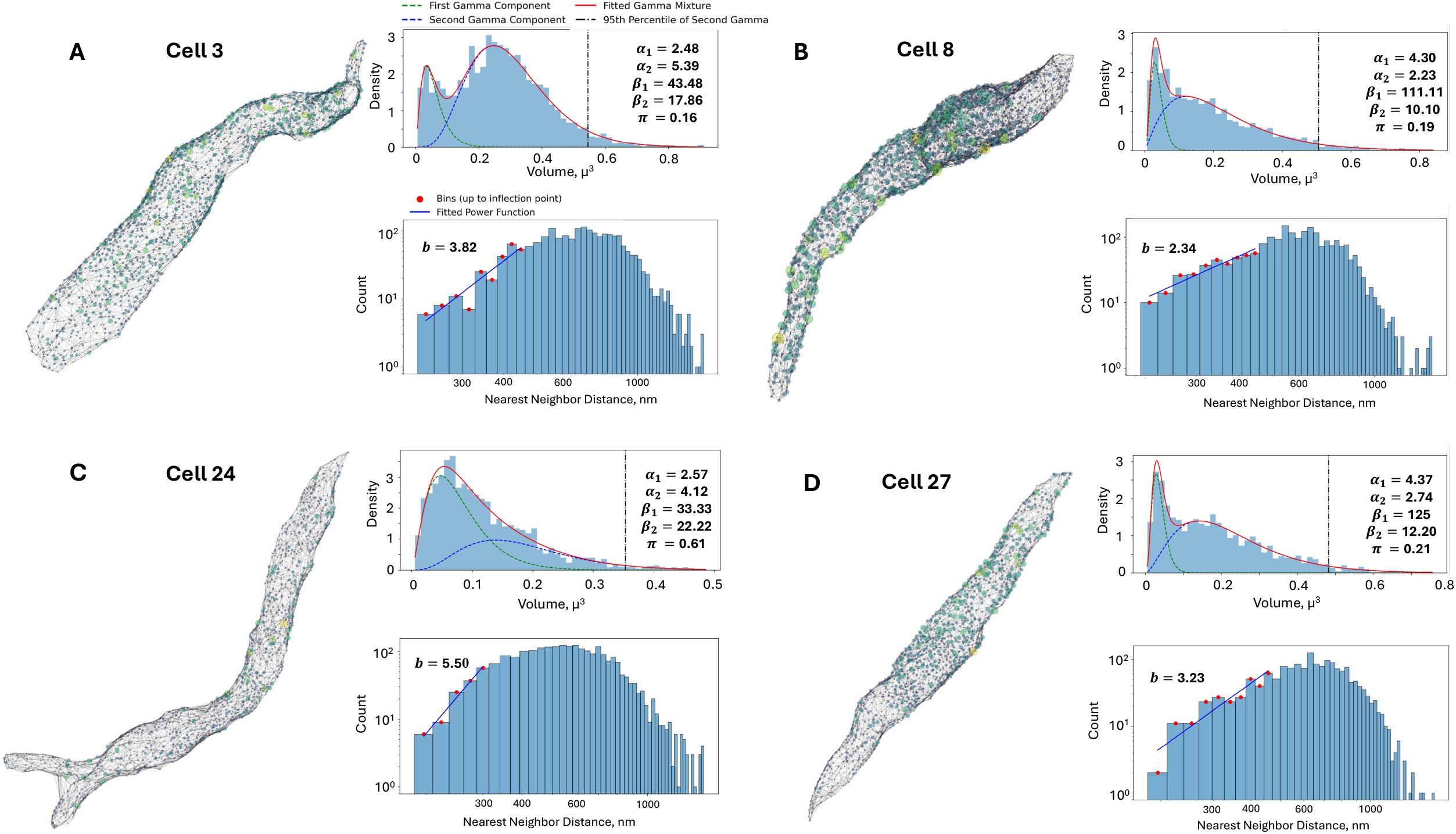
Analysis of RyR cluster distribution and nearest neighbor distances in representative rabbit SANC. (**A-D**): four representative cells (3, 8, 24, and 27 in Figure 3), each presenting three key analyses. First, a 3D visualization depicts RyR clusters as colored spheres scaled to their volumes (yellow for largest, purple for smallest) within a gray mesh representing the cell surface. Second, a fitted Gamma Mixture Model for RyR cluster size distributions is shown, with blue histograms representing observed data, green and blue dashed lines indicating individual gamma components, and a red line showing the overall fitted mixture. Model parameters (α1, α2, β1, β2, *π*) and the 95th percentile of the second gamma component (black dash-dot line, marking large CRUs) are displayed. Third, a log-log histogram of RyR cluster nearest neighbor distances is presented, where blue bars show observed distances, red dots mark bins up to the inflection point used for power law fitting (blue line), and the fitted power law exponent (b) is provided. This comprehensive analysis reveals cell-to-cell variability in RyR cluster organization and spacing but overall consistency, offering insights into the spatial arrangement of Ca^2+^ release sites in SANC.

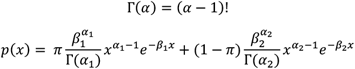

where Г(*α*) is the Gamma function, *π* is the mixture weight and *α, β* are the shape and scale parameters for their respective Gamma distribution component. The GMM was fit to the data using the *scikit-learn* library in Python, which implements the expectation-maximization algorithm to estimate the parameters of the mixture model. The number of Gamma components in the mixture was determined by evaluating models with different numbers of components and selecting the one with the lowest Bayesian Information Criterion to balance goodness of fit with model complexity.

To initialize the GMM, an estimate of the peak locations in the data was obtained using the method of moments for initial parameter estimation, which splits the sorted data at the median and uses the statistics of each half to approximate the parameters of two gamma distributions. The two most prominent peaks were identified and used as initial estimates for the means of the Gamma components. The optimization of the GMM parameters was performed using the minimize function from the *scipy.optimize* module, with bounds set on the parameters to ensure they remain positive and the mixing coefficient remains between 0 and 1. The GMM analysis revealed that the distribution of RyR cluster sizes in the rabbit sinoatrial node cells is best described by a mixture of two Gamma components with results for individual cells are shown in Table 2.

After application of Tukey’s Fences on the sizes, the RyR clusters were aggregated into a set of 70,849 clusters. Next a histogram with 100 bins was created and a GMM was fit on top of this aggregated set. The fitted parameters of the GMM were as follows in Figure 5: *α*_1_ = 4.35, *β*_1_ = 125, *α*_2_ = 2.22, *β*_2_ = 10.64, *π* = 0.16. The first gamma component had an average of 0.0333 μm^3^ and a standard deviation of 0.0160 μm^3^, while the second gamma component had an average of 0.2083 μm^3^ and a standard deviation of 0.1398 μm^3^. These results provide a comprehensive characterization of the RyR cluster size distribution in the rabbit SANC, highlighting the presence of two distinct subpopulations with different average sizes and variability. The kurtosis of the first and second Gamma components was calculated to be 1.38 and 2.70, respectively. The higher kurtosis of the second component suggests that it represents a subpopulation of larger RyR clusters with a heavier tail in the distribution. The 95th percentile of the second Gamma component was found to be 0.478 μm^3^, which was used as a threshold to define the tail of the distribution representing the largest RyR clusters used in further analysis.

**Figure 5.**
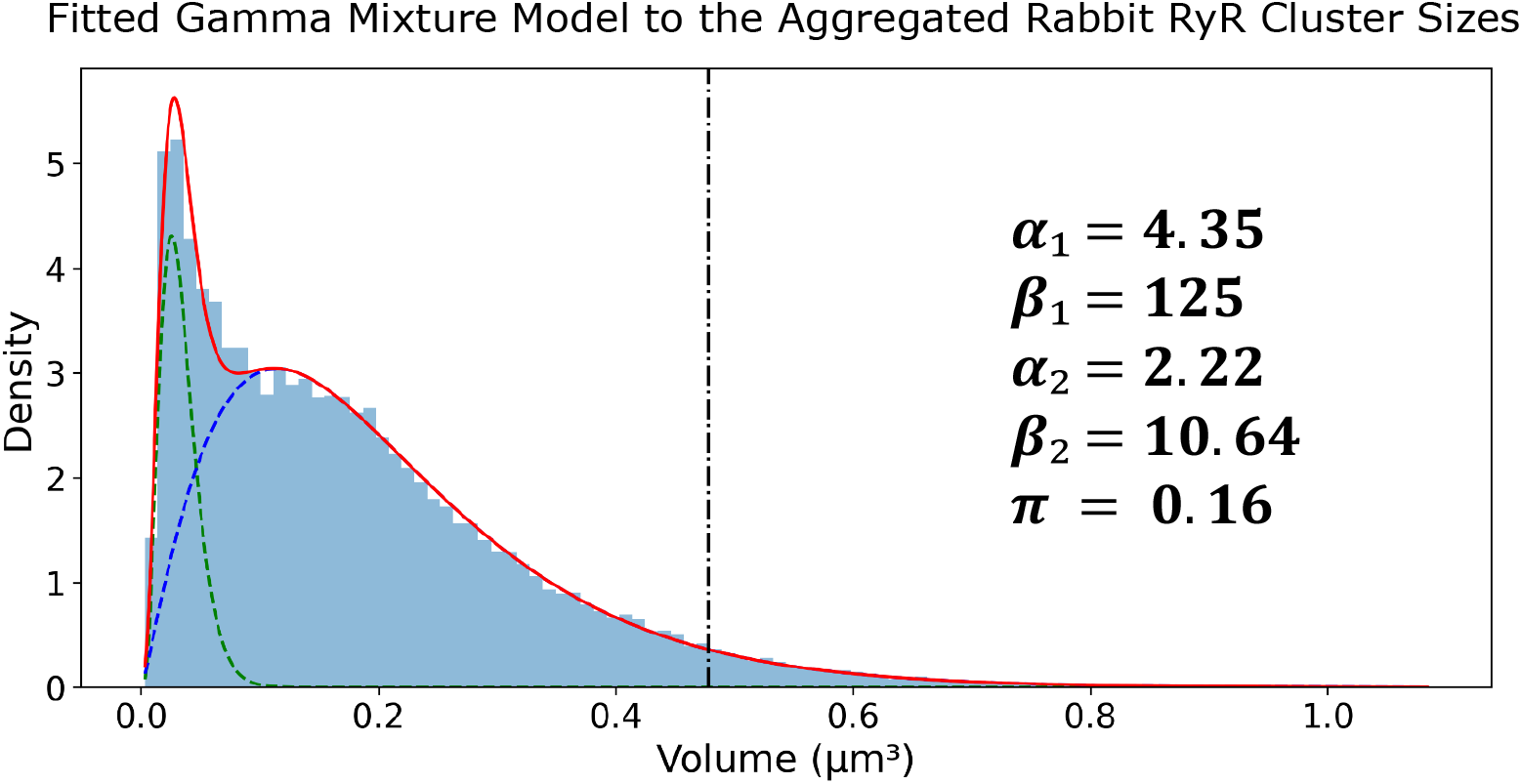
Gamma mixture model analysis of RyR cluster sizes in rabbit SANC. Probability density function (PDF) of RyR cluster sizes derived from 31 rabbit SANC, encompassing 70,982 clusters. The distribution is fitted with a two-component Gamma Mixture Model, with parameters (α1, β1, α2, β2, *π*) displayed. The blue histogram represents experimental data, while green and blue dashed lines show individual gamma components. The red line depicts the overall fitted mixture, and the black dash-dot line marks the 95th percentile of the second gamma component (marking large CRUs).

### 3.2. Repulsive behavior of RyR clusters in spatial distribution

To investigate the spatial arrangement of RyR clusters, the aggregated distances between nearest neighbor clusters were analyzed. The distribution of nearest neighbor distances provides insights into potential spatial interactions, such as repulsion or attraction, among the clusters. The overall mean distance between nearest neighbor RyR clusters was found to be 638.62 nm, with a standard deviation of 208.14 nm (Table 1). Histograms of the nearest neighbor distances for each individual SANC were constructed using 50 bins and a power law function which was fit to the ascending part of the distribution up to the inflection point, determined as the bin where the histogram values first fell below half of the peak value.

The power law function used to model the nearest neighbor distances is given by: *p*(*x*) = *Ax*^*b*^, where *p*(*x*) is the probability of observing a nearest neighbor distance, *A* is a normalization constant, and *b* is the power law exponent. Transforming the dataset into a log-log plot, a linear relationship is obtained between *log*(*p*(*x*)) and *log*(*x*). This transformation allows for a more straightforward analysis of the power law behavior. The resulting equation in log-log form is: *log*(*p*(*x*)) = *log*(*A*) + *b* · *log*(*x*). The variable *b* characterizes the spatial distribution of the clusters, with *b* ≈ 1 indicating a random Poisson process, *b* > 1 suggesting repulsion, and *b* < 1 indicating clustering. The fitting of the power law function to the ascending part of the histogram until the inflection. Four example cells and their associated analysis are shown in Figure 4. Results in Figure 6B and Table 2 show that each cell exhibited repulsion with *b* > 1. Figure 7 is a visual representation of the spatial distribution of RyR clusters, showing varying levels of cluster density on the surface mesh on sample cell 14. Higher density areas are colored in blue→green→yellow, representing RyR cluster “hotspot zones”, while low density areas are in purple which exhibit strong repulsive forces.

**Figure 6.**
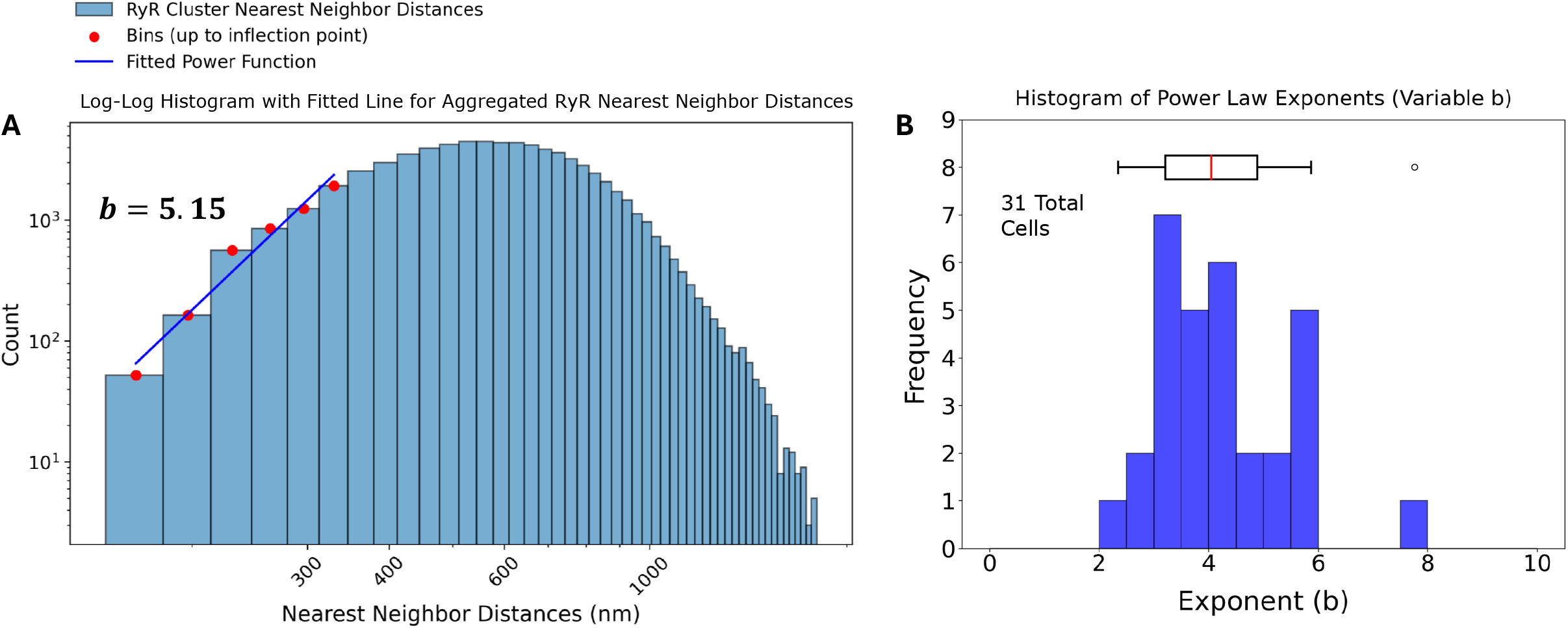
Analysis of RyR cluster nearest neighbor distances in rabbit sinoatrial node cells. (A) Log-log histogram of nearest neighbor distances (NND) for ryanodine receptor (RyR) clusters. The blue bars represent the observed NND distribution. Red dots indicate bins up to the inflection point, which were used to fit a power function (blue line). The fitted power law exponent b = 3.81 is displayed, suggesting strong repulsion among RyR clusters at short distances. (B) Histogram of power law exponents (b) derived from individual analysis of 31 cells, with an accompanying box plot.

**Figure 7.**
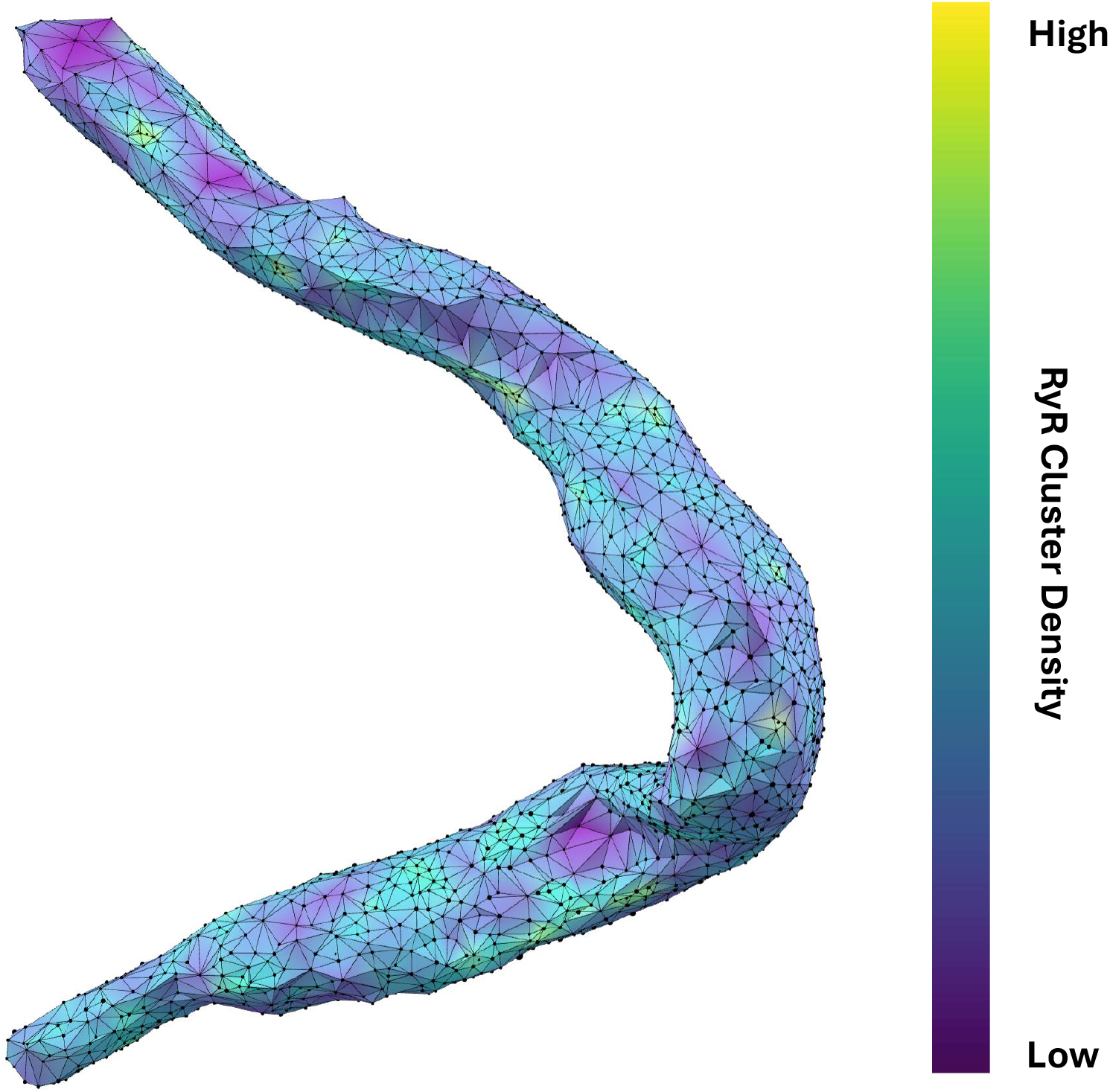
Spatial distribution and connectivity analysis of RyR clusters in a rabbit SANC. Detailed visualization of RyR cluster distribution in cell #14 (in Figure 3). The cell’s shape is outlined by a network of interconnected black vertices, each representing an RyR cluster. The surface coloring, transitioning from purple to yellow (*via* blue and green), indicates the density and connectivity of these clusters. Purple areas signify regions where clusters strongly repel each other, resulting in lower cluster density and fewer neighboring connections. Conversely, blue→green→yellow areas represent regions where this repulsion is weaker, leading to higher density regions with more interconnected clusters. The varying degrees of repulsion between RyR clusters across different areas of the cell provide insights into the spatial organization of Ca^2+^ release sites, which influence CICR and AP firing rate. Specifically, this visualization technique effectively highlights potential hotspots of Ca^2+^ signaling activity within the cell that is likely associated with higher density of RyR clusters.

After application of Tukey’s Fences on the distances, the aggregated set of 70,887 nearest neighbor distances was also analyzed and the results of the analysis revealed a power law exponent of b = 5.15, also indicating a strong repulsion among the RyR clusters at short distances (Figure 6A).

### 3.4. Structure-function relationship of CRU network in numerical simulations of SANC function

We simulated AP firing in numerical SANC models (Figure A1) with different distributions of sizes and locations. With respect to sizes, we tested and compared two distributions: one was the realistic (i.e. heterogeneous) distribution obtained in SIM data adopted to the SANC model (re-scaled and re-binned) and the other was the distribution featuring identical CRU sizes, with each CRU size fixed to 48 RyRs representing the average CRU size in the realistic distribution. It is also important to note, that each cell model had the same total number of RyRs. With respect to CRU locations, we tested three CRU distributions of NNDs: (i) uniformly random, (ii) realistic, and iii) crystal-like grid (Figure A2). The distributions were generated by our CRU repulsion algorithm that dynamically changed positions of CRUs from uniformly random locations with no repulsion towards a crystal-like structure with the strongest repulsion. The change was performed in 100 repulsion steps (see details in Appendix and Videos S1 and S2). The realistic NND distribution was found at an intermediate repulsion step with the repulsion, yielding an average NND value closely matching to that measured experimentally.

Finally, each model was tested in basal state AP firing and in the presence of β-adrenergic receptor (βAR) stimulation. Thus, we tested operation of 6 different SANC models under two conditions, resulting in 12 simulations, altogether. The results of these simulations are summarized in Figure 8. The simulations (see examples in Videos S3-S8) showed that for each CRU location distribution type (random, realistic, crystal) the models with realistic CRU sizes had a faster AP firing rate vs. those with identical CRU sizes. At the same time, the absolute values of AP firing rates were the largest (reflecting shortest cycle length) in cells with random CRU locations (Figure 8A), but the smallest (reflecting longest cycle length) in crystal-like grid with the strongest CRU repulsion (Figure 8C). The scenario with realistic locations (Figure 8B) had an intermediate AP firing rate of 157 bpm, close to the experimentally reported rates of isolated rabbit SANC.

**Figure 8.**
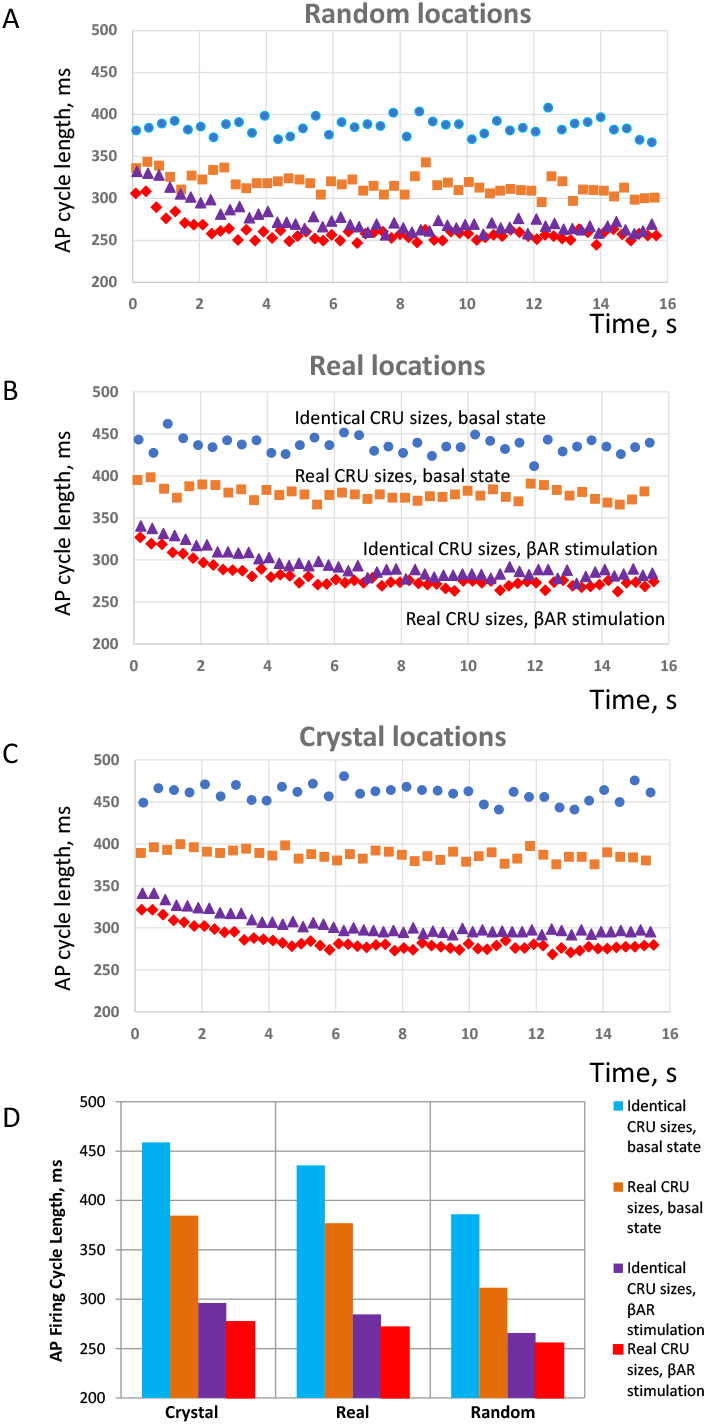
**Results of numerical model simulations of SANC function: heterogeneities in both CRU sizes and locations facilitate LCR propagation and increase AP firing rate in a cooperative manner but decrease effect of βAR stimulation in terms of relative change in AP firing rate; at the same time, the presence of heterogeneities in both sizes and locations allows to reach higher absolute AP firing rates during βAR stimulation. (A-C):** Intervalograms for AP cycle length (CL) in 12 numerical model simulations during 16 s for 6 models featuring different distribution of CRU sizes and distances in basal state and during βAR stimulation (shown by text labels in the panel B). (**D**): Graph showing average CL calculated for steady-state AP firing during 6 to 16 s of the simulations (see also Videos S3-S8).

With respect to βAR stimulation, its effect in terms of *relative* change of AP firing CL (or rate) increased as the order of CRU sizes and position increased, i.e. heterogeneity or disorder decreased, and reached its maximum in the model with identical CRU located in crystal-like grid with the least heterogeneity and maximum order. See panel D, blue and magenta bars in “Crystal section”, 459 vs. 296 ms, yielding a 55% increase in AP firing rate from 131 vs 202 bpm. And vice versa, βAR stimulation effect decreased as the disorder of CRU sizes and position increased (i.e. order decreased) and reached its minimum in the model with realistic CRU located in (uniformly) random locations. See Figure 8D, orange and red bars in “Random” section, 311 vs. 255 ms, yielding only a 22% increase in AP firing rate from 192 vs 225 bpm. On the other hand, the disorder (heterogeneity) of CRU sizes and locations allowed the models to reach higher absolute values of AP firing rates (shorter cycle lengths) in both basal state and during βAR stimulation. As the result, the highest beating rate of 225 bpm among all 12 tested scenarios was achieved in the SANC model with the highest degree of disorder representing a cooperative effect of the most heterogeneous (realistic) sizes and uniformly random locations of their CRUs (red diamonds in panel C). The model with realistic CRU sizes and locations demonstrated a 39% βAR increase in AP firing rate (377 vs 271 ms of CL, or 159 vs. 220 bpm).

Next, we analyzed our simulations to get insights into the biophysical mechanism of the contribution of heterogenous CRU network to regulate pacemaker function. According to our previous studies, the critical part of the diastolic depolarization is the AP ignition [16] at the threshold of L-type Ca^2+^ current (I_CaL_) activation. In our model each CRU size is given by the number of RyRs embedded in that CRU. Therefore, in all 12 simulation scenarios, we determined the total number of RyRs in all firing CRUs at -45 mV, i.e. at the threshold of I_CaL_ activation (Cav1.3 current component in our model of I_CaL_). We found that the AP cycle length in basal state and during βAR stimulation was linked to that number of firing RyRs (Figure 9), reflecting the degree of order and disorder (heterogeneity) of CRUs and their ability to self-organize to fire earlier and synchronously within the AP cycle.

**Figure 9.**
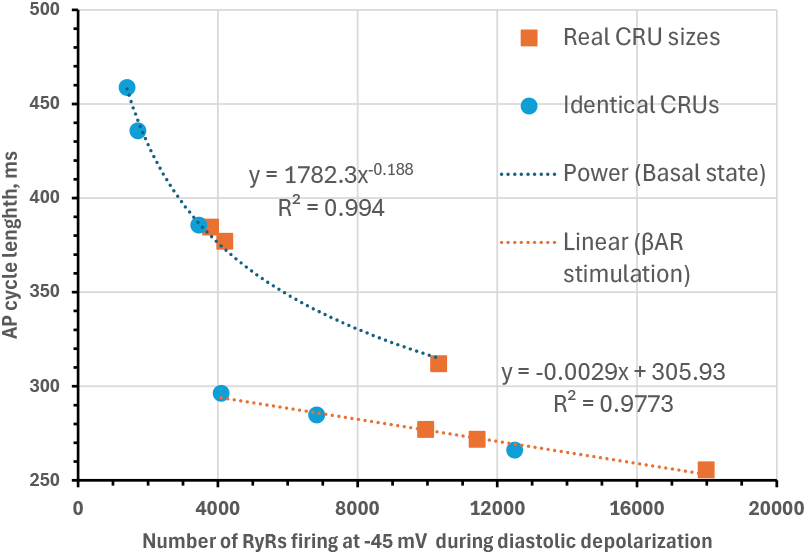
Biophysical mechanism of how CRU distribution affects AP firing rate. Shown is the relationship between of AP cycle length (CL) vs. the total number of RyRs in all firing CRUs at -45 mV, i.e. at the threshold of I_CaL_ activation. In both basal state firing (fitted by power function, R^2^=0.994) and during βAR stimulation (fitted by linear function, R^2^=0.9773) CL was closely linked to the RyR firing at this critical timing within diastolic depolarization.

## 4. Discussion

### 4.1. Result Summary: major findings

The present study, for the first time, examined the detailed structure of RyR network under the cell membrane in SANC (Figure 3) at the nanoscale level allowed by SIM super-resolution microscopy. Previous lower resolution confocal microscopy studies showed that RyR appear in clusters which are mainly located under cell membrane in central (primary) SANC [17]. It was also shown that RyR-mediated LCR events, play a crucial role in the coupled clock system that ensure robust and flexible pacemaking [7]. Many important properties of the network, however, remained unknown. The major findings of the present study included:

1. Two distinct RyR cluster subpopulations with sizes adhering to a Gamma mixture distribution
2. Notable repulsion between neighboring RyR clusters
3. Model simulations showed that heterogeneities in both CRU sizes and locations facilitate CICR and increase AP firing rate in a cooperative manner but decrease effects of βAR stimulation. However, the highest possible absolute rate during βAR stimulation is reached with heterogeneities in both CRU sizes and locations.

### 4.2. New image analysis methods and numerical models

We developed a novel, high-throughput algorithm for the segmentation of peripheral RyR clusters in 3D in SANC, enabling the analysis of over 70,000 RyR clusters across 31 rabbit SANC (Figure 1). Our advanced image preprocessing techniques included 3D median filtering, signal normalization, and CLAHE [30]. It enhanced the local contrast while preserving the global image structure, facilitating more accurate segmentation. For the first time in SANC, the 3D StarDist neural network [31] was trained for the segmentation of RyR clusters and accurately predicted star-convex polyhedra representations for each cluster, improving upon traditional methods that rely on watershed algorithms and often struggle with overlap. The ground truth data for training the neural network was prepared using Squassh [32] and fine-tuned with the Adaptive Watershed tool in ORS Dragonfly [36]. Further refinement through DBSCAN clustering removed false positives. To accurately capture the peripheral RyR clusters directly interacting with cell membrane ion channels and transporters critical for heart rate regulation, the cell periphery was isolated using an alpha shape algorithm, constructing a minimal bounding surface that closely conforms to the shape of the data points, effectively representing the cell’s geometry.

We also modified our previous 3D numerical model of SANC [22] by including new formulations for CRU function (see Appendix for details). Recent theoretical studies [37-39] showed that Ca^2+^ release activation in a CRU (i.e. Ca^2+^ spark activation) depends substantially on the CRU size, i.e. the number of RyRs in a CRU. Thus, our new formulations were created to reflect actual distributions obtained by segmentation of SIM images: they generate repulsion between the CRUs (Figure A2, Videos S1 and S2) and CRU of various sizes (Figure A3). Using the new formulations, we created and tested a series of models with real and extreme settings that revealed importance of heterogeneities in CRU sizes and locations and their joint regulation of CICR to ensure robust and flexible pacemaker function.

### 4.3. Structure-function relations of RyR network in SANC

While it is known that LCRs play an important role in pacemaker function [7] the detailed mechanisms regarding how LCRs emerge from the RyR network remain mainly unknown. Because the RyR network operates via CICR, its properties must critically depend on both CRU *sizes* (generating stronger releases) and *distances* that define whether the release of each CRU can reach and activate its neighbors (i.e. “when can a Ca^2+^ spark jump?” [21]). Our previous theoretical studies of CRU function in SANC showed first that synchronization of stochastic CRUs creates a rhythmic Ca^2+^ clock in SANC via repetitive phase-like transitions within a perfect rectangular grid of CRU network [19]. Further linking this model with membrane clock revealed that RyR-Na/Ca exchanger (NCX)-SERCA local crosstalk ensures pacemaker cell function at rest and during the fight-or-flight reflex [20].

While previous studies tested the effects of CRU sizes and distances in separation, here, for the first time, we found that they act together to regulate pacemaking (Figure 8). The CRUs form a functional network in which Ca^2+^ release events self-organize (via CICR) into local Ca^2+^ oscillators that create a rhythmic diastolic Ca^2+^ signal known as the “Ca clock” that together with the “membrane clock” forms a coupled clock system that ensures robust and flexible pacemaking [7]. The diastolic Ca^2+^ signal acts not alone, but within strong positive feedback mechanism (known as AP ignition [16]) with NCX, membrane potential, and I_CaL_. The self-organized Ca^2+^ signal formed by CRUs at the threshold of I_CaL_ (at the ignition point) must play a critical role in the ignition and ultimately AP firing rate regulation.

Our simulations showed that this critical signal at the threshold of I_CaL_ is tightly linked to the structure of the RyR network. The heterogeneity in both CRU sizes and distances strongly increase the CICR and the AP firing rate in both basal state and during βAR stimulation (Figure 9). Thus, to reach the maximal relative increase in firing rate and at the same time maximum absolute rate, the CRU network can transform to increase its heterogeneity, i.e. to increase the contribution of larger RyR clusters. Recent dSTORM studies in ventricular myocytes showed that the RyR clusters can expand and coalesce after application of isoproterenol [40]. Future dSTORM studies will show whether this also happens in SANC allowing the RyR network to transform itself to reach maximum performance in fight-or-flight reflex.

We have previously shown [22] that disorder in CRU locations increases CICR due to Poisson clumping characterized by empty spaces and clusters. The CRU clustering makes it easier for released Ca^2+^ to reach a neighboring CRU. But why does a CRU network with heterogeneous CRU sizes also enhance CICR and AP firing rate? Larger CRUs are naturally present in such network (Figures 4 and 5, beyond vertical line). These larger CRUs have a greater probability and capacity for Ca^2+^ release [39], acting as central hubs that initiate and amplify CICR to the surrounding smaller CRUs. This multi-scale spatial organization enhances the likelihood of CICR propagation from large clusters to smaller ones, promoting synchronization necessary for effective Ca^2+^ signaling.

Pilot dSTORM studies of RyR clusters in rat SANC (abstracts [41,42]) fully support the results of the present study. It was found that the cluster sizes vary from 10 to 150 RyRs and they organized in couplons, i.e. in apposition to L-type Ca^2+^ channels found largely at the outer membrane in SAN cells. The heterogenous RyR2 clusters were also found with often small inter-cluster distances, which is indicative of inter-cluster RyR2 activation.

### 4.4. Limitations and future studies

Our cell database was limited to 31 cells and our model simulations were limited to 12 basic scenarios. Future studies in a larger number of cells and with even stronger imaging resolution, like dSTORM (up to 20 nm resolution [43]), will clarify further details of RyR network structure and its relationship with other networks, formed e.g. by mitochondria [44] or Ca^2+^ channels of different types and isoforms [45-47] directly or indirectly interacting with RyR network.

Future numerical modeling will include and test the functional importance of these relationships. The presence of spatial repulsion between CRUs suggests that there exists a molecular mechanism that actively maintains minimum spacing. Further biophysical studies will also clarify the nature of CRU repulsion and, possibly, coalescence [40]. Thus, the CRU sizes are likely determined, in part, by the repulsion/coalescence balance controlled by autonomic modulation to reach maximal (or optimal) effect.

The antibody mAb C33-3 used to detect RyRs in SANC was generated against RyR2, but also cross-reacts with RyR1. However, using RT-PCR, it was reported that sinoatrial node contains both RyR2 and RyR3, but not RyR1 [48]. The relative abundance of RyR2 and RyR3 is distinct between cells located at the periphery vs. those at the center of the sinoatrial node: RyR2 is higher at the periphery and RyR3 higher in cells at the center of the node [49]. Thus, the present study did not investigate all RyRs but was focused on cardiac type RyR2. Further studies are needed to clarify CRU composition and function with respect to both RyR2 and RyR3 isoforms in SANC residing in different regions of the node.

### 4.5. Implications for clinical and aging research

Disorder within biological systems tends to increase with aging across all scales. Thus, heterogeneity (i.e. disorder) in CRU sizes and spatial distribution is also expected to increase in older people, but as a result, CICR will actually be enhanced, thereby elevating the AP rate. This enhancement could partially compensate for the reduced intrinsic heart rate commonly associated with aging. However, the same increase in structural disorder of the RyR network would diminish β-adrenergic responsiveness, as demonstrated in this study, contributing to (and explaining in part) the decline in heart rate reserve observed in older people.

As we mentioned in Introduction, dysfunction of SANC can lead to sick sinus syndrome associated with a variety of life-threatening arrhythmias [3-6]. Electronic pacemakers are used to treat this condition but impose lifestyle restriction and can cause severe side effects. To improve electronic pacemakers and create new approaches, for example, biological pacemakers [50], it is important to understand the natural origin of the heart-beat that is “still mysterious after all these years” (cited from [23]). SANC operates via a coupled-clock mechanism [7] and its “Ca^2+^ clock” operates via synchronous and rhythmic release of Ca^2+^ via RyR cluster network. Our new findings of structure-function relation of the network can help direct the development of new prophylactic and therapeutic strategies. Specifically, our results suggest that healthy operation of the network requires a delicate balance of order and disorder in RyR cluster positions and sizes that determines and ensures the respective healthy balance of robustness and flexibility of cardiac pacemaking. Dysfunction of this balance could be a new factor in heart rate reserve decline in aging and disease. Our numerical model provides a new platform to investigate how molecular defects in CRUs and CRU network structure may disrupt pacemaking in diseases like sick sinus syndrome. Incorporating measured CRU network properties will help to provide mechanistic insights into pacemaker pathologies and develop new biological pacemakers.

### 4.6. Conclusions

CICR interplay of CRUs of various sizes and locations regulates and optimizes cardiac pacemaker cell operation under various physiological conditions and could be a key factor in heart rate reserve decline in aging and disease. Overall, our study elucidates how the specialized architecture of the CRU network in SANC regulates ignition and propagation of Ca^2+^ sparks to generate rhythmic AP firing. The combination of super-resolution imaging and computational modeling provides a powerful approach to link subcellular Ca^2+^ release patterns to whole-cell pacemaker activity.

## 5. Appendix: Details of a novel numerical model of SANC

In the present study we investigate functional importance of both heterogeneous CRU sizes and locations (Figure A1). To get further insights into operation of the CRU network featuring realistic locations and sizes (as observed in SIM images), we developed a new numerical cell model that has a novel Ca^2+^ release mechanism that dependents on the CRU size. The new model is based on our previous model of rabbit SANC [22] that included a full set of electrophysiological equations and local Ca^2+^ dynamics generating LCRs via CRUs, but it was limited to CRUs of identical sizes. With respect to CRU location, the new model simulates repulsion between CRUs (see Videos S1 and S2) that was revealed in SIM images. The LCR emergence is approximated in the model at the scale of an individual CRU, i.e. LCRs appear as Ca^2+^ sparks and abrupted Ca^2+^ waves (wavelets, as observed in prior experimental studies) via saltatory propagation of CICR among neighboring CRUs. Each CRU, in turn is represented by a cluster of RyRs embedded in the JSR, located in close proximity (20 nm) to cell surface membrane. We did not simulate operation of each individual RyR, but (in the spirit of multi-scale modeling [51]) we tuned the CRU formulations, i.e. clusters of RyRs, by performing simulations of the Stern model of CRU [52] featuring individual RyR firing. All equations of this new numerical model (and model tuning) including novel formulation of CRU activation and generation of realistic heterogeneous CRU distributions (both size and location) are described below.

### 5.1. Cell geometry, compartments, voxels and membrane patches

We model a small SANC shaped as a cylinder of 53.28 μm in length and 6.876 μm in diameter. The cell membrane electrical capacitance of 19.8 pF is similar to that of 20 pF in Zhang et al. model [53] of a central SANC. Details of local Ca^2+^ dynamics under the cell membrane are simulated on a sub-micron resolution square grid (120 nm, i.e. similar to SIM resolution used in this study). The grid divides the cell membrane and the submembrane space (dubbed subspace) into respective membrane patches and subspace voxels. Locations within the grid are defined by coordinates in the respective plane of the cell surface cylinder: along the cylinder (*x* axis) and around the cell cross-section (*y* axis). Our cell partitioning and respective voxel sizes are schematically illustrated in Figure A1. To avoid special considerations at the cell borders, the ends of the cylinder are connected (yielding a torus). Our cell compartments and voxel structure are essentially similar to those in Stern et al. model [21] that approximates Ca^2+^ dynamics in 3 dimensions at the single RyR scale. However, we limited the cell partition to only three nested layers of voxels of substantially different scales reflecting respective essential Ca^2+^ cycling components and processes happening at these scales (described below).

#### 5.1.1. Submembrane voxels

The first, most detailed level of approximation of local Ca^2+^ dynamics is achieved via 79920 voxels, 444 in x and 180 in y. Each voxel of 120×120×20 nm (ΔxΔyΔr) is located under the entire cell membrane, including the 20 nm dyadic space (or cleft space) separating CRUs and the cell membrane. This thin layer of voxels describes local Ca^2+^ release from individual CRUs, CRU-to-CRU interactions via Ca^2+^ diffusion and CRU interactions with cell membrane (including I_CaL_, I_NCX_, I_CaT_, and I_bCa_). Ca^2+^ currents were computed for each membrane patch (120×120 nm) to generate local Ca^2+^ fluxes contributing to local Ca^2+^ dynamics.

#### 5.1.2. JSR level voxels (“ring” voxels)

The next approximation level of Ca^2+^ dynamics is a deeper layer of voxels that have a larger size of 360×360×800 nm (*ΔxΔyΔr*) that includes the scale of JSR depth (60 nm). Each ring voxel has its cytoplasmic part and FSR part and some ring voxels are diffusively connected to JSRs (Figure A1). While it appears that the geometric scale of ring voxel is of an order of magnitude larger than the JSR size, the actual FSR volume within each ring voxel is comparable with the JSR volume (described below in details). Thus, this level of voxels describes the local dynamics of Ca^2+^ transfer from FSR to JSR as well as enhanced local Ca^2+^ pumping and diffusion fluxes due to close proximity to Ca^2+^ release and Ca^2+^ influx in its neighboring submembrane voxels.

#### 5.1.3. The core

The rest of the cell does not have further geometric partitioning and it is lumped to one compartment “the core”, where local Ca^2+^ dynamics is less important. The core also has its cytosolic and FSR parts (as in ring voxels). Thus, it describes the bulk Ca^2+^ uptake from cytosol to FSR and further accumulation, diffusion, and redistribution of the pumped and released Ca^2+^ within the cell interior.

#### 5.1.4. JSR

While individual release channels are not modelled directly, each CRU is characterized by its number of RyR channels determining its JSR size. Because the thickness of pancake-like JSR is fixed to 60 nm, the volume of each JSR is directly proportional to the number of RyRs (*N*_*RyR*_) residing in the JSR and facing the dyadic space. Because RyRs form a 30×30 nm crystal grid and because the dyadic space (as a part of submembrane space) split into voxels of 120×120 nm in xy plane, each xy voxel size can accommodate 120*120/(30*30) =16 RyRs. This dictates possible *N*_*RyR*_ and respective geometries that JSRs can have: *N*_*RyR*_ varies as multiples of 16, i.e. *N*_*RyR*_ =16*n, where is n is positive integer, i.e. 16, 32, 48, 64, etc. Thus, each CRU is characterized by a set of adjacent submembrane voxels fully populated by RyRs and our computer algorithm configured CRU geometries as ideal squares of the voxels, when possible (e.g. 2×2 for 4 voxels populated by 64 RyRs or 3×3 for 9 voxels populated by 144 RyRs) or form incomplete squares for intermediate cases with random rotation.

Based on available experimental data, our previous SANC model [21] had 196,000 RyRs. That cell model had length of 112 μm and diameter of 8 μm, yielding cell cylinder surface area of *π*\*8*112 = 2,814.86701762 μm^2^, with RyR density of 196,000/2,814.86701762 = 69.6302876026 ∼70 RyRs per μm^2^. A similar RyR density was also used in our original CRU-based model of SANC [22] and in the present model. We examine here two types of JSR size models: 1) all JSRs have identical number of RyRs, i.e. homogeneous JSR models; 2) JSR sizes are distributed as in experimental data. For a correct comparison of structure-function performance, we generate all models with approximately the same number (83,009) of RyRs per cell determined by the aforementioned density of 70 RyRs per μm^2^ and total cell surface area in our cell model of 2\**π*\* 3.495043* 54 = 1,185.84015258 μm^2^. The exact number of RyRs per cell slightly varies in different RyR distributions because (i) RyRs populate JSRs as multiples of 16 as described above and (ii) there is a small remainder when CRUs are populated following a specific RyR distribution. When we created CRUs and populated them with RyRs in the cell model using experimental data, we first calculated the number of RyRs for CRU size (16, 32, 48, etc) by simple scaling: ModelCRUSizeDistr(i) = 83,009* ExpCRUSizeDistr(i) /Σ ExpCRUSizeDistr(i).

In our original model [22] with identical CRUs we simulated various degrees of order and disorder in JSR positions by generating respective various degrees of deviations from their crystalline positions. In our new model we developed a different approach that simulates CRU repulsion found in SIM images of RyR clusters. In short, the new algorithm considers CRUs as charged particles of same polarity with the effective charge and mass proportional to their sizes (i.e. *N*_*RyR*_). Initially CRUs are distributed randomly (at time 0). JSR overlaps are excluded, i.e. any two JSRs cannot occupy the same cell volume. Then CRUs move freely (but with friction) driven by respective Coulomb forces. The friction is important for the system to ultimately (in ∼100 steps) evolve into a steady state exhibiting a crystal-like structure due to strong repulsion (see Videos S1 and S2, for CRUs of heterogeneous and identical sizes, respectively). Thus, iteration steps create a series of distributions of CRU positions reflecting various degrees of repulsion (or order and disorder) (Figure A2). It is important to note, however, that we don’t know the nature of repulsion in real CRU distributions, and we found all parameters in the repulsion algorithm empirically. We use Coulomb forces here just to create a computationally effective repulsion process that generates respective CRU distributions with various degrees of repulsion. Those distributions are further tested to evaluate the functional effect of the repulsion per se, regardless the nature of the repulsion.

Each JSR is diffusely linked to the network of FSR that uptakes Ca^2+^ from cytosol via sarco/endoplasmic reticulum Ca-ATPase (SERCA) pumping. We place JSRs inside the respective ring voxels just below their outer side facing the cell membrane (Figure A1). Volumes of the ring voxels are kept the same by their extending into the core by the exact volume that the JSR occupies at their outer side. Thus, the actual core volume is calculated as the volume of the cylinder core decreased by the volume of all JSR volumes.

### 5.2. Ca^2+^ cycling

#### 5.2.1. Free SR (FSR)

As mentioned above, each ring voxel and the core are further partitioned into cytosol and FSR fractions. We model FSR as homogeneously distributed network within the cytosol. The cytosol fraction is set to 0.46 and FSR fraction to 0.035 [21]. The remainder presumably contains myofilaments, mitochondria, nucleus, and other organelles. In turn, each FSR portion has capability to pump Ca^2+^ locally from the respective cytosol portion of the same voxel, simulating local SERCA function. Because the submembrane voxels are extremely thin, only 20 nm depth, their contribution to Ca^2+^ pumping and intra-SR diffusion are negligible and not modelled.

#### 5.2.2. SR Ca^2+^ pump

The SERCA pump is present uniformly throughout the cell, transferring Ca^2+^ from the cytosolic to the FSR compartment of each voxel (of ring and core) with Ca^2+^ uptake rate given by the reversible Ca^2+^ pump formulation adopted from Shannon et al. [54]

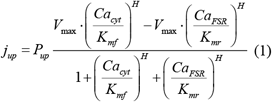

where *P*_*up*_ = 0.012 mM/ms, *K*_*mf*_ = 0.000246 mM, *K*_*mr*_ = 1.7 mM, and H = 1.787.

#### 5.2.3. Ca^2+^ diffusion within and among cell compartments

Ca diffusion fluxes between voxels within and among cell compartments are approximated by the first Fick’s law:

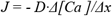

where *D* is a diffusion coefficient, and *Δ[Ca]/Δx* is Ca^2+^ concentration gradients, i.e. *Δ[Ca]* is the concentration difference and *Δx* is the distance between the voxel centers.

The respective rate of change of [Ca] is defined as

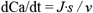

where *s* is the diffusion area sharing by voxels and *v* is the receiving volume. Thus,

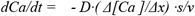

For any two diffusively interacting voxels with volumes *v*_*1*_ and *v*_*2*_, Ca^2+^ dynamics is described by a set of differential equations:

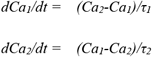

where *Ca*_*1*_ and *Ca*_*2*_ are Ca^2+^ concentrations in the respective voxels and

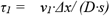

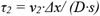

*τ*_*1*_ (or *τ*_*2*_, symmetrically) is the respective time constant of *Ca*_*1*_ change in time in a special case if the other compartment with volume *v*_*2*_ is substantially larger than *v*_*1*_ (i.e. *v*_*2*_ *>> v*_*1*_) and therefore *Ca*_*2*_ remains approximately constant. In general case, the set is analytically solved to the respective exponential decays:

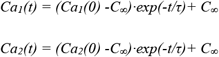

where *C*_*∞*_*= (Ca*_*1*_*(0) ·v1 + Ca*_*2*_*(0) · v*_*2*_*)/(v*_*1*_ *+ v*_*2*_) is equilibrium concentration (t = *∞*) in both voxels defined by the matter conservation principle and *τ =τ*_*1*_*·τ*_*2*_*/(τ*_*1*_ *+ τ*_*2*_) is the common time constant of the exponential decay of the system to reach the equilibrium. The respective Ca^2+^ change from its initial value in voxel *v*_*1*_ over time is given as follows:

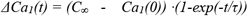

Then, by substituting C_*∞*_ we get:

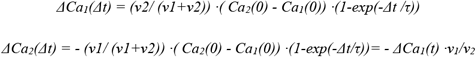

These formulations are used in all our computations of [Ca] changes for all neighboring voxels within and among cell compartments for the model integration for each time update *Δt* (during time tick or several time ticks for slower processes). In our computer algorithm we calculate the fractional Ca^2+^ change (*FCC*) before the model run and use it simply as a scaling factor to determine actual [Ca] change from the difference in [Ca] between any two diffusely interacting voxels at the beginning of each integration step (from *t=0* to *t=Δt*). Thus,

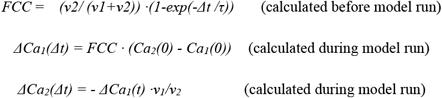

In the case of identical voxels, i.e. within subspace and within ring (i.e. when *v*_*1*_ *= v*_*2*_ and *τ*_*1*_*= τ*_*2*_) the formulations are simplified to:

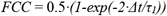

Note 1: If *Δ*t/*τ*_*1*_ << 1 then *FCC* can be approximated (e.g. via respective Taylor series) as

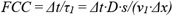

Further, if *v*_*1*_ can be described as *v*_*1*_ *= s · Δx*, e.g. for diffusion along the cell length (axis *x* in our model) *FCC* can be further simplified to

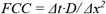

However, because *FCC* is calculated only once, before the model run, and does not carry any additional computational burden during actual simulations of Ca^2+^ dynamics, we always use here the full approximation for the diffusion, i.e. more precise exponential decay, rather than a linear change over *Δt*. An advantage of this approach is that the model features more stable behavior in case we want to vary cell geometry, cell compartments, voxel sizes, or integration time (*Δt*).

Note 2: We have only a fraction of cell volume occupied with cytosol (or FSR). However, the same fraction will be for *v* and *s* in τ formulations, and it cancels. The volume ratios *v*_*2*_*/(v*_*1*_*+v*_*2*_) and *v*_*1*_*/v*_*2*_ remain also unchanged because the fraction factor also cancels. Thus, all above formulations with formal geometric volumes are also valid for fractional volumes, assuming that the fraction of cytosol (or FSR) is evenly distributed within the volume.

#### 5.2.4. Junctional SR (JSR) and Ca^2+^ diffusion between JSR and FSR

Each JSR is refilled with Ca^2+^ locally from FSR network via a fixed diffusional resistance yielding a mono-exponential process with a fixed time constant tau. In our original model [22] with fixed JSR sizes (144 RyRs) this tau was set to 40 ms similar to previous common pool models that approximated diffusion between JSR and FSR pools [55]. Here we want to do the same for JSRs of various sizes. We do not consider Ca^2+^ diffusion within the JSR (i.e. each JSR Ca^2+^ is described by one variable), but each JSR can touch several FSR voxels (Figure A1). As described above, complex JSR geometries represent a collection of *N* adjacent 120×120×60 nm parts/blocks, with each part accommodating 16 RyRs (*N= N*_*RyR*_*/16*). Hence, we set a diffusional link for each such JSR part to its adjacent FSR voxel (360×360 nm in xy). The net Ca^2+^ flux to a given JSR from FSR is calculated as a sum of Ca^2+^ fluxes via those individual links, with each link transferring Ca^2+^ flux with *τ* = N*·40 ms · v*_*FSR*_*/(v*_*FSR*_ *+ v*_*JSR*_), where *v*_*JSR*_ is volume of a given JSR and *v*_*FSR*_ is FSR volume within a ring voxel.

#### 5.2.5. Spark activation and inactivation

Each CRU can be either in open or closed state. The capability of a given CRU to open, i.e. to generate a Ca^2+^ spark, is controlled by its JSR Ca^2+^ loading. Experimental and theoretical studies showed that sparks are generated when SR Ca^2+^ loading reaches a certain critical level [39,56]. This critical *Ca*_*jSR*_ level is implemented in our mechanism of spark activation by prohibiting CRU firing while SR Ca^2+^ loading remains below 300 μM (*CaSRfire*). When JSR is refilled with Ca^2+^ above this level, it can open. The switch from close state to open state is probabilistic. The probability density for a given closed CRU to open is described by a power function of Ca^2+^ concentration (*Ca*) in the dyadic space. In our original model [22] CRU size was fixed to 144 RyRs. The probability *p*_*144*_ for CRU of this size to open during a short time interval *TimeTick* was given by

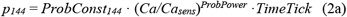

where *Ca*_*sens*_ = 0.15 μM was sensitivity of CRU to Ca^2+^ in dyadic space, *ProbConst*_*144*_ *=*0.00027 ms^-1^ was open probability rate at Ca^2+^ = *Ca*_*sens*_, and *ProbPower* = 3 defining the cooperativity of CRU activation by Ca^2+^ in dyadic space. In our new model, CRU sizes vary and therefore we adopted the original formulation as follows:

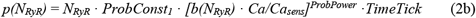

In the new formulation we scaled the probability rate with respect to *N*_*RyR*_ in the CRU, so that *ProbConst*_*1*_ *= ProbConst*_*144*_*/144 = 1.875*10*^*-6*^ ms^-1^. We also introduced a boundary factor *b(N*_*RyR*_) that reflects a lower probability to ignite a spark by an RyR opening in smaller CRUs due to stronger escape of the released Ca^2+^ to bulk cytosol from the tight dyadic space via closer CRU boundary to the open channel. This Ca^2+^ escape results in a lower (smaller) spatial Ca^2+^ profile from one open RyR, resulting in a weaker CICR with respect to neighboring RyRs and lower probability of spark ignition. To find the boundary factor *b(N*_*RyR*_), we performed a series of numerical model simulations using Stern at al. spark model [52] that describes individual RyRs within a CRU. Results of these simulations are presented in Figure A3.

Our previous studies showed that the critical interaction of RyRs within a CRU with respect to CICR and spark activation is described by interaction profile, *ψ(r)*, i.e. the profile of Ca^2+^ drop in dyadic space when one RyR stays opens, and especially Ca^2+^ at the nearest RyR neighbor [39,57]. Thus, we found a series of interaction profiles *ψ(r)* as a function of *N*_*RyR*_ for a large variety of CRU sizes from 3×3 to 15×15 RyRs. Then we plotted Ca^2+^ at the nearest RyR neighbor of the open channel and fitted it to a phenomenological exponential curve *Ca*_*neighbor*_*(N)* that closely described waning RyR interactions for smaller CRUs (grey line in Figure A3C). To get the boundary factor in Equation 2b, we normalized the exponential curve to its value at 144 (i.e. *b(144)=1*) to be consistent with our previous formulation for the CRU with 144 RyRs [22]. Because *Ca*_*neighbor*_*(144)* = 25.387 μM, we finally get

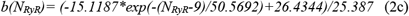

Each time tick our computer algorithm tries to activate a closed CRU by generating a random number within (0,1). If this number less than *p* (in Equation 2b), then the CRU opens. The Ca^2+^ current amplitude, *I*_*spark*_, is defined by spark activation kinetics *a(t)*, RyR unitary current (*I*_*RyR,1mM*_, the current via a single RyR at 1 mM of Ca^2+^ gradient), *N*_*RyR*_, and concentration difference between inside and outside JSR:

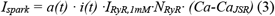

*I*_*RyR,1mM*_ is set to 0.35 pA[52]. Spark activation *a(t)* kinetics is described as a single exponential time-dependent process with *τ*_*a*_ = 10 ms:

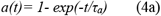

Because each open RyR undergoes time-dependent inactivation, this is also reflected in our CRU model by adding simple inactivation kinetic variable *i(t)* with a time constant *τ*_*i*_ *= 1/k*_*om*_ *=* 8.547 ms (inactivation rate constant *k*_*om*_ = 0.117 ms^-1^ in Stern et al. spark model [52]):

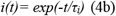

#### 5.2.6 Spark termination mechanism

Our spark termination mechanism is based on the current knowledge in this research area [52,57,58], i.e. a Ca^2+^ spark is generated via CICR among individual RyRs within a CRU and it sharply terminates due to induction decay [58], CICR interruption [52], or a phase transition (similar to that known in Ising model)[57] when RyR current *I*_*RyR*_*(t)* becomes too small (due to JSR Ca^2+^ depletion) to further support the CICR among RyRs. The specific value of the critical current *I*_*spark_termination*_ is defined by interactions of individual RyRs and beyond the capability of our CRU-based model. Thus, we terminated spark when the spark current *I*_*spark*_*(t)* becomes very small, equal to RyR current *I*_*RyR*_*(t)*, i.e. the current via one RyR channel at a given JSR Ca^2+^ load.

#### 5.2.7 Ca^2+^ buffering

Cytosolic Ca^2+^ is buffered by calmodulin (0.045 mM) throughout the cell: submembrane voxels, ring voxels, and the core. Each JSR features Ca^2+^ buffering with calsequestrin (30 mM).

#### 5.2.8 Summary of equations of local Ca^2+^ dynamics

In a submembrane voxel:

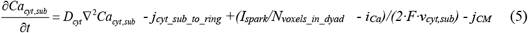

*I*_*spark*_ in voxels outside dyadic space is absent. Each CRU releases Ca^2+^ (given by *I*_*spark*_) into its dyadic space. *I*_*spark*_ is evenly distributed among submembrane voxels of the dyadic space (*N*_*voxels_in_dyad*_). *I*_*spark*_ is given in Equation 3 and it is positive, i.e. increasing [Ca] in the submembrane voxel. *i*_*Ca*_ is the sum of local Ca^2+^ transmembrane currents (described in details below in Electrophysiology section) via the membrane patch facing this submembrane voxel:

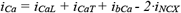

*I*_*CaL*_ is included in this equation only for submembrane voxels facing a CRU (*I*_*CaL*_ is injected into the subspace voxels of the respective dyadic space). The local Ca^2+^ currents *i*_*CaL*_, *i*_*CaT*_, *i*_*bCa*_ have inward direction and (by convention) are defined as negative. Therefore, the minus sign before *i*_*Ca*_ in Equation 5 ensures positive change in [Ca] in the submembrane voxel by respective Ca^2+^ influx. During diastole local *i*_*NCX*_ also flows inwardly, but NCX exchanges 1 Ca^2+^ ion to 3 Na ions. Hence, the inward *i*_*NCX*_ generates a Ca^2+^ efflux. That is why it has a different sign.

In a voxel of ring layer:

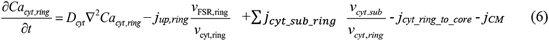

∑ *j*_*cyt_sub_ring*_ is the sum of diffusion fluxes from neighboring smaller submembrane voxels

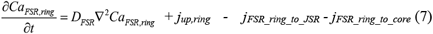

The SERCA uptake flux *j*_*up*_ is given by Equation 1.

In a given JSR:

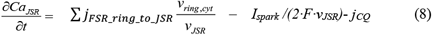

where *I*_*spark*_ is given by Equation 3.

∑ *j*_*FSR_ring_to_JSR*_ is the sum of diffusion fluxes between JSR and respective FSR parts of the neighboring ring voxels. *j*_*CQ*_ is Ca^2+^ flux of Ca^2+^ buffering by calsequestrin:

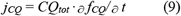

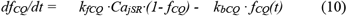

In the core:

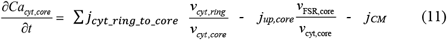

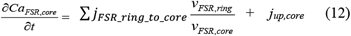

where ∑ *j*_*cyt_ring_to_core*_ and ∑ *j*_*FSR_ring_to_core*_ are the respective sums of diffusion fluxes in cytosol and FSR of all ring voxels.

In any cytoplasmic voxel (subspace, ring, and the core):

*j*_*CM*_ is Ca^2+^ flux of Ca^2+^ buffering by calmodulin:

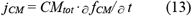

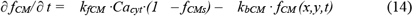

### 5.3. Electrophysiology

Major changes of the model include introduction of local Ca^2+^ currents and modulation of local currents by local Ca. To introduce local currents and local modulation by Ca, the cell membrane is partitioned into small patches, with each patch facing its respective subspace voxel. We also omitted sustained inward current *I*_*st*_ and background Na current *I*_*bNa*_ because thus far the molecular identities for these currents have not been found and these currents are likely produced by NCX or other currents [7].

#### 5.3.1. Fixed ion concentrations, mM

*Ca*_*o*_ *= 2*: Extracellular Ca^2+^ concentration.

*K*_*o*_ *= 5.4*: Extracellular K concentration.

*K*_*i*_ *=140*: Intracellular K concentration.

*Na*_*o*_ *= 140*: Extracellular Na concentration.

*Na*_*i*_ *=10*: Intracellular Na concentration.

#### 5.3.2. The Nernst equation and electric potentials, mV

*E*_Na_ = *E*_T_ · ln(Na_o_/Na_i_): Equilibrium potential for Na

*E*_*K*_ *= E*_*T*_ *· ln(K*_*o*_*/K*_*i*_): Equilibrium potential for K

E_Ks_ = E_T_ · ln[(K_o_ + 0.12· Na_o_)/(K_i_ + 0.12 · Na_i_)]: Reversal potential of I_Ks_

Where *E*_*T*_ is “*RT/F*” factor = 26.72655 mV at 37°C,

*E*_*CaL*_ *= 45*: Apparent reversal potential of *I*_*CaL*_

*E*_*CaT*_ *= 45*: Apparent reversal potential of *I*_*CaT*_

#### 5.3.3. Membrane potential, V_m_

Net membrane current determines time derivative of the membrane potential.

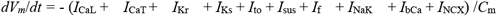

#### 5.3.4. Formulation of cell membrane ion currents

Kinetics of ion currents are described by gating variables (described below) in respective differential equations

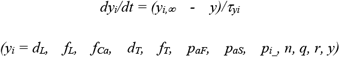

*τ*_*yi*_: time constant for a gating variable *y*_*i*_.

*α*_*yi*_ and *β*_*yi*_: opening and closing rates for channel gating.

*y*_*i*_,_*∞*_: steady-state curve for a gating variable *y*_*i*_.

L-type Ca^2+^ current *(I*_*CaL*_)

The whole cell *I*_*CaL*_ is calculated as a sum of local currents *i*_*CaL,i*_ in each dyadic space.

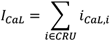

This reflects reports that L type Ca^2+^ cannels (LCCs) are colocalized with RyRs [59]. Thus, the whole cell maximum *I*_*CaL*_ conductance (*g*_*CaL*_) is distributed locally among dyadic spaces proportionally to their areas:

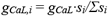

where *s*_*i*_ is submembrane area of a *i-th* CRU and *Σs*_*i*_ is the area occupied by all CRUs. The respective local *i*_*CaL,i*_ is calculated in each CRU as a sum of two currents generated by two major LCC isoforms Ca_v1.2_ and Ca_v1.3_ present in SANC [2,60]

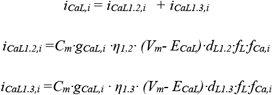

where *η*_*1.2*_ and *η*_*1.3*_ are respective fractions of Ca_v1.2_ and Ca_v1.3_ isoforms (*η*_*1.2*_ *+ η*_*1.3*_ = 1). Ca_v1.3_ has a lower (more hyperpolarized) voltage activation threshold.

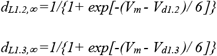

We set *V*_*d1.2*_ the midpoint of *I*_*CaL1.2*_ activation to -6.6 mV as in earlier SANC models of Wilders et al. [61] and Dokos et al. [62]. This value of *V*_*d1.2*_ is relatively high with respect to other SANC models and therefore was considered in this study (and our previous study [22]) to simulate *I*_*CaL*_ generated by Ca_v1.2_. *V*_*d1.3*_ the midpoint of *I*_*CaL1.3*_ activation is set to -13.5 mV. The resultant shift of activation increased *d*_*L1.3*,_*∞* vs. *d*_*L1.2*,_*∞* within the range of diastolic depolarization by a factor of ∼3 [22], generating a much larger current and stronger recruitment of CRUs to fire during diastolic depolarization, consistent with the current concept of important role of Ca_v1.3_ isoform in cardiac pacemaker function [46,47]. Other parameters of L-type Ca^2+^ currents are the same for both isoforms:

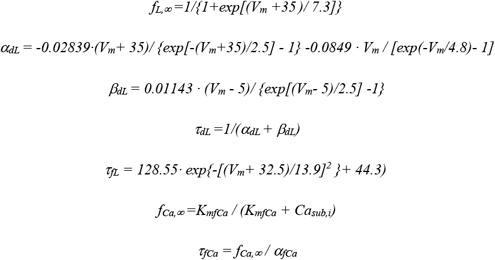

Ca-dependent *I*_*CaL*_ inactivation is described by parameters *K*_mfCa_ and *α*_fCa_.

T-type Ca^2+^ current *(I*_*CaT*_)

It is based on formulations of Demir et al.[63] and modified by Kurata et al.[55].

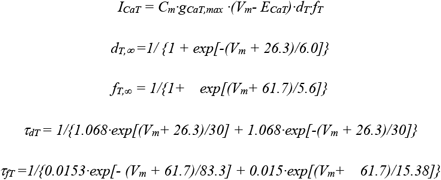

The whole cell *I*_*CaT*_ is evenly distributed over cell membrane patches to generate respective homogeneous Ca^2+^ influx. In each submembrane *i-th* voxel *I*_*CaT,i*_ *= I*_*CaT*_*/N*_*voxels*_

Rapidly activating delayed rectifier K^+^ current (*I*_Kr_)

It is based on formulations of Zhang et al. [53], further modified by Kurata et al. [55].

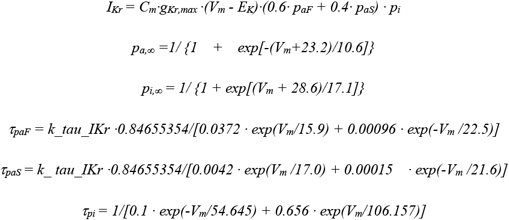

Slowly activating delayed rectifier K^+^ current (*I*_Ks_)

It is based on formulations of Zhang et al. [53].

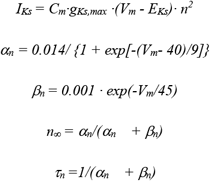

4-aminopyridine-sensitive currents (*I*_4AP_ =*I*_to_ *+ I*_sus_)

It is based on formulations of Zhang et al. [53].

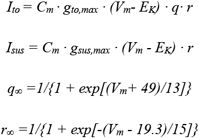

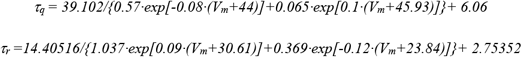

Hyperpolarization-activated, “funny” current (*I*_*f*_)

It is based on formulations of Wilders et al.[61] and Kurata et al.[55].

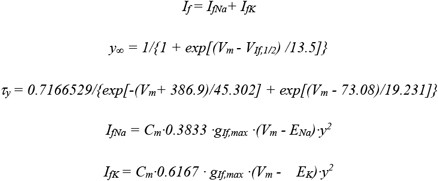

Na^+^-K^+^ pump current (*I*_*NaK*_)

It is based on formulations of Kurata et al.[55], which were in turn based on the experimental work of Sakai et al.[64] for rabbit SANC.

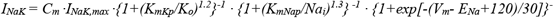

Ca-background current (*I*_*bCa*_)

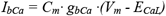

The whole cell *I*_*bCa*_ is evenly distributed over cell membrane to generate respective homogeneous Ca^2+^ influx. In each submembrane *i-th* voxel *I*_*bCa,i*_ *= I*_*bCa*_*/N*_*voxels*_

Na-Ca exchanger current (*I*_NCX_)

It is based on original formulations from Dokos et al. [62]. *I*_*NCX*_ is modulated by local Ca^2+^ and therefore the whole cell *I*_*NCX*_ is calculated as a sum of local currents *I*_*NCX,i*_ in respective membrane patches facing each submembrane voxel. Thus,

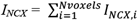

For membrane voltage *V*_*m*_ in each *i*-th membrane patch with subspace *Ca*_*sub,i*_, the respective local NCX current (*I*_*NCX,i*_) is calculated as follows:

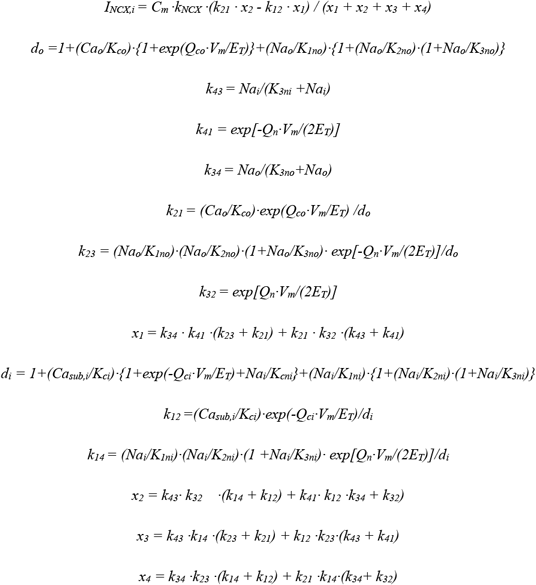

### 5.4. Initial values

#### 5.4.1. Electrophysiology

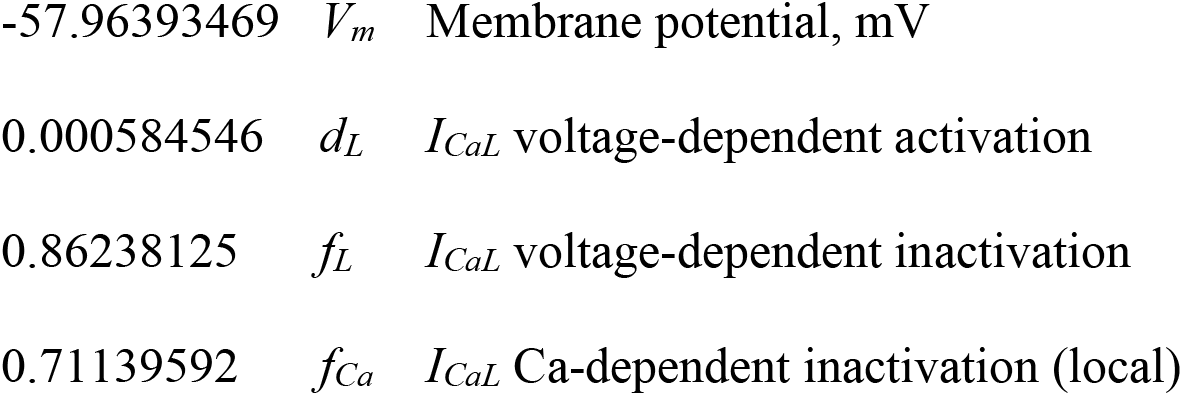
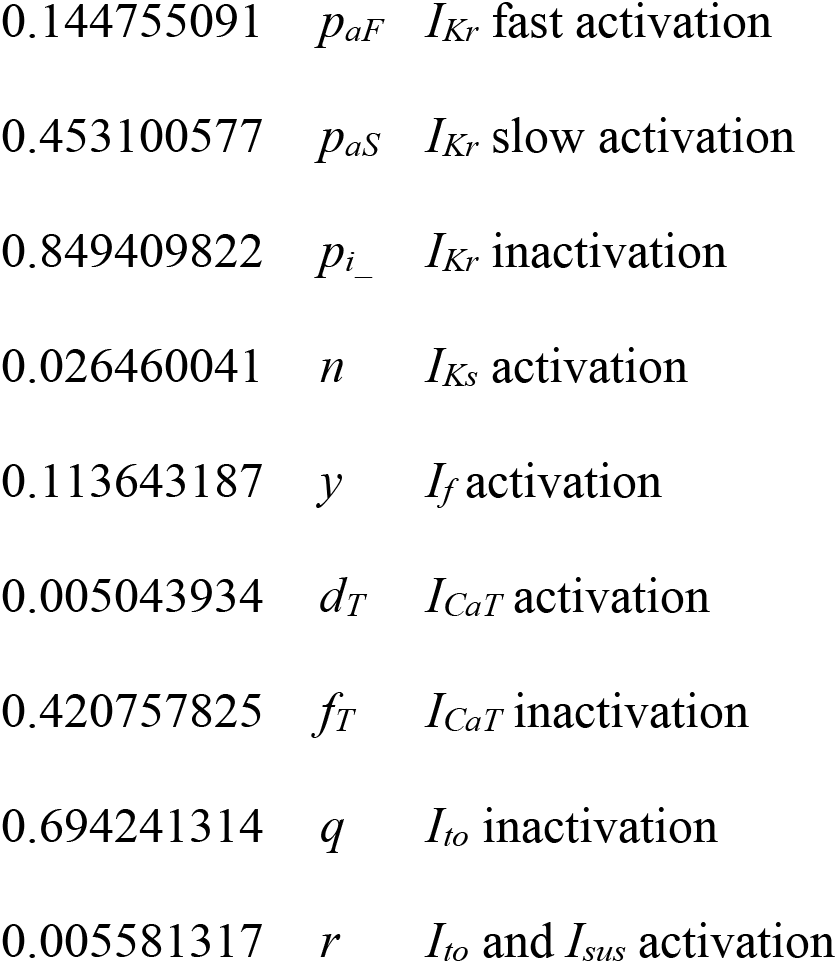

#### 5.4.2. Ca^2+^ dynamics

200 nM [Ca] in cytosol

1 mM [Ca] in FSR

0.8 mM Ca concentration in JSR

0.0787 Fractional occupancy of calmodulin by Ca^2+^ in cytoplasm

0.6 Fractional occupancy of calsequestrin by Ca^2+^ in JSR

All CRUs are set in the closed state

### 5.5. Simulations of βAR stimulation effect

Effect of βAR stimulation is modelled essentially as we previously reported [22] by increasing *I*_*CaL*_, *I*_*Kr*_, *I*_*f*_, and Ca^2+^ uptake rate by FSR via SERCA pumping. Specifically whole cell maximum *I*_*CaL*_ conductance *g*_*CaL*_ is increased by a factor of 1.75 from 0.464 to 0.812 nS/pF; whole cell maximum *I*_*Kr*_ conductance *g*_*Kr*_ is increased by a factor of 1.5 from 0.05679781 to 0.085196715 nS/pF; the midpoint of *I*_*f*_ activation curve is shifted to more depolarized potential by 7.8 mV from -64 mV to -56.2 mV; and the maximum Ca^2+^ uptake rate *P*_*up*_ is increased by a factor of 2 from 0.012 mM/ms to 0.024 mM/ms.

### 5.6. Model Integration

The model code was written in Delphi Language (Delphi 10.4) and was computed with a fixed time tick of 0.0075 ms on a workstation running Windows 10 with Intel® Xeon® W-2145 CPU @3.7GHz processor.

### 5.7. Appendix Glossary and Parameters’ Values

Fixed ion concentrations

*Ca*_*o*_=2 mM: Extracellular [Ca]

*K*_*o*_=5.4 mM: Extracellular [K]

*Na*_*o*_=140 mM: Extracellular [Na]

*K*_*i*_=140 mM: Intracellular [K]

*Na*_*i*_=10 mM: Intracellular [Na]

Membrane currents

*E*_CaL_ = 45 mV: Apparent reversal potential of *I*_*CaL*_

*g*_*CaL*_ = 0.464 nS/pF: Conductance of *I*_*CaL*_

*η*_*1.2*_ *= η*_*1.3*_ = 0.5: contributions of Ca_v1.2_ and Ca_v1.3_ isoforms to total g_*CaL*_

*K*_mfCa_ = 0.03 mM: Dissociation constant of Ca^2+^ -dependent *I*_CaL_ inactivation

*beta*_fCa_ = 60 mM^-1^ · ms^-1^: Ca^2+^ association rate constant for *I*_*CaL*_.

*alfa*_fCa_ = 0.021 ms^-1^: Ca^2+^ dissociation rate constant for *I*_caL_

*E*_CaT_ = 45: Apparent reversal potential of *I*_*CaT*_, mV

*g*_*CaT*_ = 0.1832 nS/pF: Conductance of *I*_*caT*_

*g*_*If*_ = 0.105 nS/pF: Conductance of *I*_*f*_

*V*_*If,1/2*_ = -64: half activation voltage of *I*_f_, mV

*g*_*Kr*_ = 0.05679781 nS/pF: Conductance of delayed rectifier K current rapid component

*k_ tau_IKr* = 0.3: scaling factor for *tau_paF* and *tau_paS* used to tune the model

*g*_*Ks*_ = 0.0259 nS/pF: Conductance of delayed rectifier K current slow component

*g*_*to*_ = 0.252 nS/pF: Conductance of 4-aminopyridine sensitive transient K^+^ current

*g*_sus_ = 0.02 nS/pF: Conductance of 4-aminopyridine sensitive sustained K^+^ current

*I*_NaKmax_ = 1.44 pA/pF: Maximum Na^+^/K^+^ pump current

*K*_mKp_ = 1.4 mM: Half-maximal *K*_o_ for *I*_NaK_.

*K*_mNap_ = 14 mM: Half-maximal *Na*_i_ for *I*_NaK_.

*g*_bCa_ = 0.003 nS/pF: Conductance of background Ca^2+^ current,

*k*_NCX_ = 48.75 pA/pF: Maximumal amplitude of *I*_*NCX*_

Dissociation constants for NCX

*K*_1ni_ = 395.3 mM: intracellular Na binding to first site on NCX

*K*_2ni_ = 2.289 mM: intracellular Na binding to second site on NCX

*K*_3ni_ = 26.44 mM: intracellular Na binding to third site on NCX

*K*_1no_ = 1628 mM: extracellular Na binding to first site on NCX

*K*_2no_ = 561.4 mM: extracellular Na binding to second site on NCX

*K*_3no_ = 4.663 mM: extracellular Na binding to third site on NCX

*K*_ci_ = 0.0207 mM: intracellular Ca^2+^ binding to NCX transporter

*K*_co_ = 3.663 mM: extracellular Ca^2+^ binding to NCX transporter

*K*_cni_ = 26.44 mM: intracellular Na and Ca^2+^ simultaneous binding to NCX

NCX fractional charge movement

*Q*_ci_= 0.1369: intracellular Ca^2+^ occlusion reaction of NCX

*Q*_co_=0: extracellular Ca^2+^ occlusion reaction of NCX

*Q*_n_= 0.4315: Na occlusion reactions of NCX

Ca buffering

*k*_*bCM*_ = 0.542 ms^-1^: Ca^2+^ dissociation constant for calmodulin

*k*_*fCM*_ = 227.7 mM^-1^· ms^-1^: Ca^2+^ association constant for calmodulin

*k*_*bCQ*_ = 0.445 ms^-1^: Ca^2+^ dissociation constant for calsequestrin

*k*_*fCQ*_ = 0.534 mM^-1^· ms^-1^: Ca^2+^ association constant for calsequestrin

*CQ*_*tot*_ = 30 mM: Total calsequestrin concentration

*CM*_*tot*_ = 0.045 mM: Total calmodulin concentration

SR Ca^2+^ ATPase function

*K*_*mf*_ = 0.000246 mM: the cytosolic side *Kd* of SR Ca^2+^ pump

*K*_*mr*_ = 1.7 mM: the lumenal side *Kd* of SR Ca^2+^ pump

H = 1.787: cooperativity of SR Ca^2+^ pump

*P*_*up*_ = 0.012 mM/ms: Maximal rate of Ca^2+^ uptake by SR Ca^2+^ pump

CRU (Ca release and JSR)

*Ca*_*sens*_ = 0.00015 mM: sensitivity of Ca^2+^ release to Casub

*ProbConst*_*1*_ *= 1.875*10*^*-6*^ ms^-1^: CRU probability constant per each RyR in the CRU

*ProbPower* = 3: Cooperativity of CRU activation by Casub

CaJSR_spark_activation = 0.3 mM: critical JSR Ca^2+^ loading to generate a spark (CRU can open)

*τ*_*CRUactivation*_ = 10 ms: Time constant of spark activation (*a*)

*τ*_*CRUinactivation*_ = 8.547 ms: Time constant of spark inactivation (*i*)

Iryr_at_1mM_CaJSR = 0.35 pA: unitary RyR current at 1 mM delta Ca

RyR_to_RyR_distance_um = 0.03 μm: RyR crystal grid size in JSR

JSR_depth_um = 0.06 μm: JSR depth

JSR_Xsize_um = 0.36 μm: JSR size in x

JSR_Ysize_um = 0.36 μm: JSR size in y

JSR_to_JSR_X_um = 1.44 μm: JSR crystal grid size in x

JSR_to_JSR_Y_um = 1.44 μm: JSR crystal grid size in y

Cell geometry, compartments, and voxels

*L*_cell_ = 53.28 μm: Cell length

*r*_cell_ = 3.437747 μm: Cell radius

*C*_*m*_ = 19.80142 pF: membrane electrical capacitance of our cell model with 0.0172059397937 pF/μm^2^ specific membrane capacitance calculated from 2002 Kurata et al. model [55] for its 32 pF cylinder cell of 70 μm length and 4 μm radius.

0.12 μm: the grid size

0.12 μm: Submembrane voxel size in *x*

0.12 μm: Submembrane voxel size in *y*

0.02 μm: Submembrane voxel size depth

0.36 μm: Ring voxel size in *x*

0.36 μm: Ring voxel size in *y*

0.8 μm: Ring voxel depth

0.46: Fractional volume of cytosol

0.035: Fractional volume of FSR

Ca diffusion

*Dcyt* = 0.35 μm^2^/ms: Diffusion coefficient of free Ca^2+^ in cytosol

*D*_*FSR*_ = 0.06 μm^2^/ms: Diffusion coefficient of free Ca^2+^ in FSR

## Supplementary Materials

Videos S1 and S2: The sequence of CRU distributions (for real CRU sizes and identical CRUs, respectively) generated by our CRU repulsion algorithm that dynamically changed positions of CRUs from uniformly random locations with no repulsion towards a crystal-like structure with the strongest repulsion (cylinder cell surfaces were unwrapped to squares). The change was performed in 100 repulsion steps. The CRUs (green objects) on the top panel are shown together with respective histogram of NNDs and plots of average NND and standard deviation at each step. For better representation of nonlinear changes, the first 13 steps are shown at a slower rate of 0.5s/frame, with rest steps at 0.1 s/frame. Videos S3 - S8: Ca^2+^ dynamics under the cell membrane in numerical simulations in the basal state of our 6 models with different CRU distributions as follows: S3, random positions and realistic sizes; S4, realistic positions and sizes, S5, crystal-like positions and realistic sizes; S6, random positions and identical sizes (48 RyRs); S7, realistic positions and identical sizes, S8, crystal-like positions and identical sizes. Ca^2+^ concentration is coded by red shades from black (0.15 μM) to pure red (>10 μM). CRU functional states are coded by colors as follows: CRUs in refractory are in blue, CRUs ready to release are in green, CRUs releasing Ca^2+^ are in grey shades, reflecting JSR Ca^2+^ changes, with a saturation level set at 0.3 mM. Simulation time and membrane potential are shown in the left top corner of each video.

## Author Contributions

Conceptualization, Alexander V.M., E.G.L. and V.A.M.; methodology, Alexander V.M., .V.V., O.M., M.J., S.T., Anna V.M., and V.A.M.; software, Alexander V.M., V.V., S.R., and V.A.M.; validation, Anna V.M., S.T., M.D.S., and V.A.M.; investigation, Alexander V.M., V.V., O.M., M.J., P.A.W., S.R., Anna V.M., and V.A.M.; resources, E.G.L. and M.D.S.; data curation, Alexander V.M., .V.V., and V.A.M.; formal analysis, Alexander V.M., V.V., O.M., P.A.W, S.R., and V.A.M.; writing—original draft preparation, Alexander V.M., and V.A.M.; writing—review and editing, Alexander V.M., V.V., Anna V.M., E.G.L. and V.A.M.; visualization, Alexander V.M., V.V., and V.A.M.; supervision, M.D.S., E.G.L., and V.A.M.; project administration, E.G.L., and M.D.S.; All authors have read and agreed to the published version of the manuscript.

## Funding

This research was supported by the Intramural Research Program of the National Institutes of Health, National Institute on Aging. Anna V. Maltsev acknowledges the support of the Royal Society University Research Fellowship UF160569. Valeria Ventura acknowledges the support of U-RISE undergraduate scholarship program (at the University of Maryland Baltimore County) funded by the National Institute of General and Medical Sciences.

### Institutional Review Board Statement

The animal study was reviewed and approved by the Animal Care and Use Committee of the National Institutes of Health (protocol #457-LCS-2024).

### Informed Consent Statement

Not applicable.

## Data Availability Statement

All original data files were deposited to Harvard Dataverse at https://doi.org/10.7910/DVN/N0OLGI. Code Availability: Code for image data analysis was uploaded to GitHub: https://github.com/alexmaltsev/SANC, https://github.com/valventura/SANC. The original code of the numerical model (in Delphi programing language) was published as a supplement in our last publication [22].

## Acknowledgments

The authors thank Dr. Hari Shroff (currently at Janelia Research Campus) for the opportunity to perform SIM imaging in his research laboratory at the National Institute of Biomedical Imaging and Bioengineering, NIH. The authors wish to acknowledge the assistance of Bruce D. Ziman for skillful isolation of sinoatrial node cells. The authors also thank Daniel Riordon and Larissa Maltseva for their assistance in performing SIM imaging

## Abbreviations

SANC: sinoatrial node cell
AP: action potential
LCR: local Ca^2+^ release
βAR: β-adrenergic receptor
RyR: Ryanodine Receptor
SERCA: sarco/endoplasmic reticulum Ca-ATPase
Cycle length: action potential cycle length
CRU: Ca^2+^ release unit
SR: sarcoplasmic reticulum
JSR: the junctional SR
CICR: Ca-induced-Ca release
SIM: structured illumination microscopy
NND: nearest neighbor distances
PBS: phosphate buffered saline
CLAHE: contrast-limited adaptive histogram equalization
IoU: Intersection over Union
DBSCAN: Density-Based Spatial Clustering of Applications with Noise
MAD: median absolute deviation
GMM: Gamma Mixture Model
CDF: cumulative distribution function

## FIGURE LEGENDS

**Figure A1.**
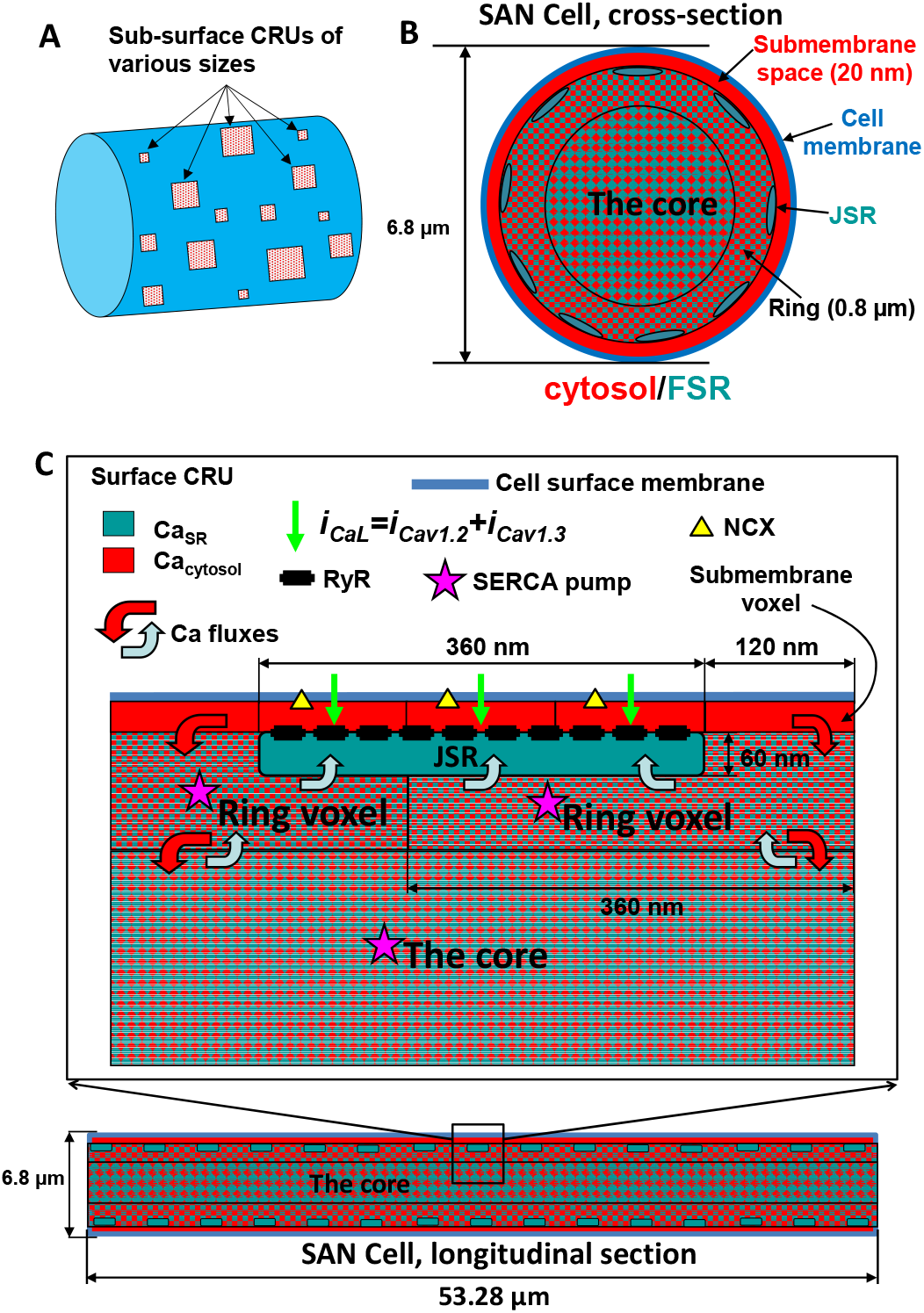
Schematic illustration of approximation of local Ca_2+_ dynamics in our CRU-based SANC model. Panel A shows the spatial positioning of Ca^2+^ release units (CRUs) beneath the cell membrane. CRUs are shown in different sizes (a key feature of our new model), reflecting different number of ryanodine receptors (RyRs) included in each CRU. In Panels B and C, a cross-sectional and longitudinal section of our simulation are presented respectively, revealing a three-layer voxel-based approximation of intracellular Ca^2+^ dynamics. These voxel layers offer a discretized 3D view of the cellular space, enabling the detailed spatiotemporal representation of Ca^2+^ signals within the cell. Specifically, the first layer (red) approximates the submembrane space where the CRUs are located and interact with cell membrane proteins. The second layer (ring) approximates cell space associated with the junctional sarcoplasmic reticulum (JSR), and third layer (the core) is representative of the bulk cytosolic space. Both the ring and the core includes network sarcoplasmic reticulum the primary intracellular Ca^2+^ storage site that is equipped with Ca^2+^ pump (SERCA, labeled by star). Collectively, this updated 3D SANC model, with its refined representation of CRU sizes and locations, provides a more accurate and realistic approximation of Ca^2+^ signaling in sinoatrial node cells. Adopted from [22].

**Figure A2.**
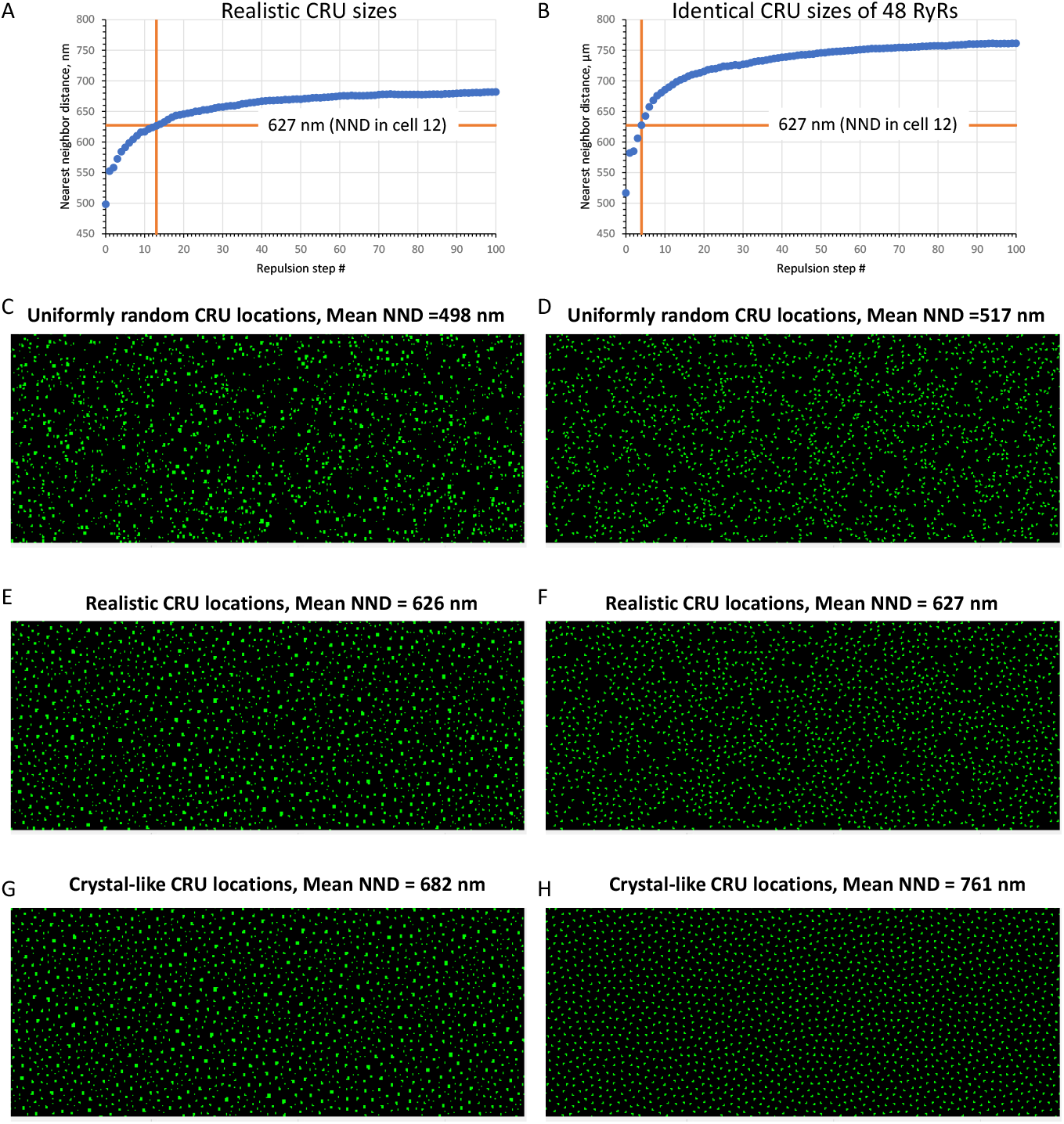
Various degrees of CRU repulsion (i.e. order and disorder) generated by our repulsion algorithm in a cell with realistic CRU sizes (as in cell #8, Table 1) (left panels) and in a virtual cell with CRUs having identical sizes accommodating 48 RyRs (right panels). Top panels (A and B) show how average NND changed in the respective CRU networks under cell membrane as repulsion transformed uniformly randomly distributed CRUs (step 0) towards crystal-like structure (step 100). An intermediate CRU network with mean nearest neighbor distance (NND) close to that measured experimentally was used in our model simulations (steps 13 and 4 for networks of realistic and identical CRU sizes, respectively, vertical orange lines in A and B). Panels C-H show CRU distributions under the cell membrane (cylinder surfaces unwrapping to squares) that were used in our numerical model simulations of SANC function. See also Videos S1 and S2 for more details, including NND distributions, mean NND, and NND standard deviation at each repulsion step.

**Figure A3.**
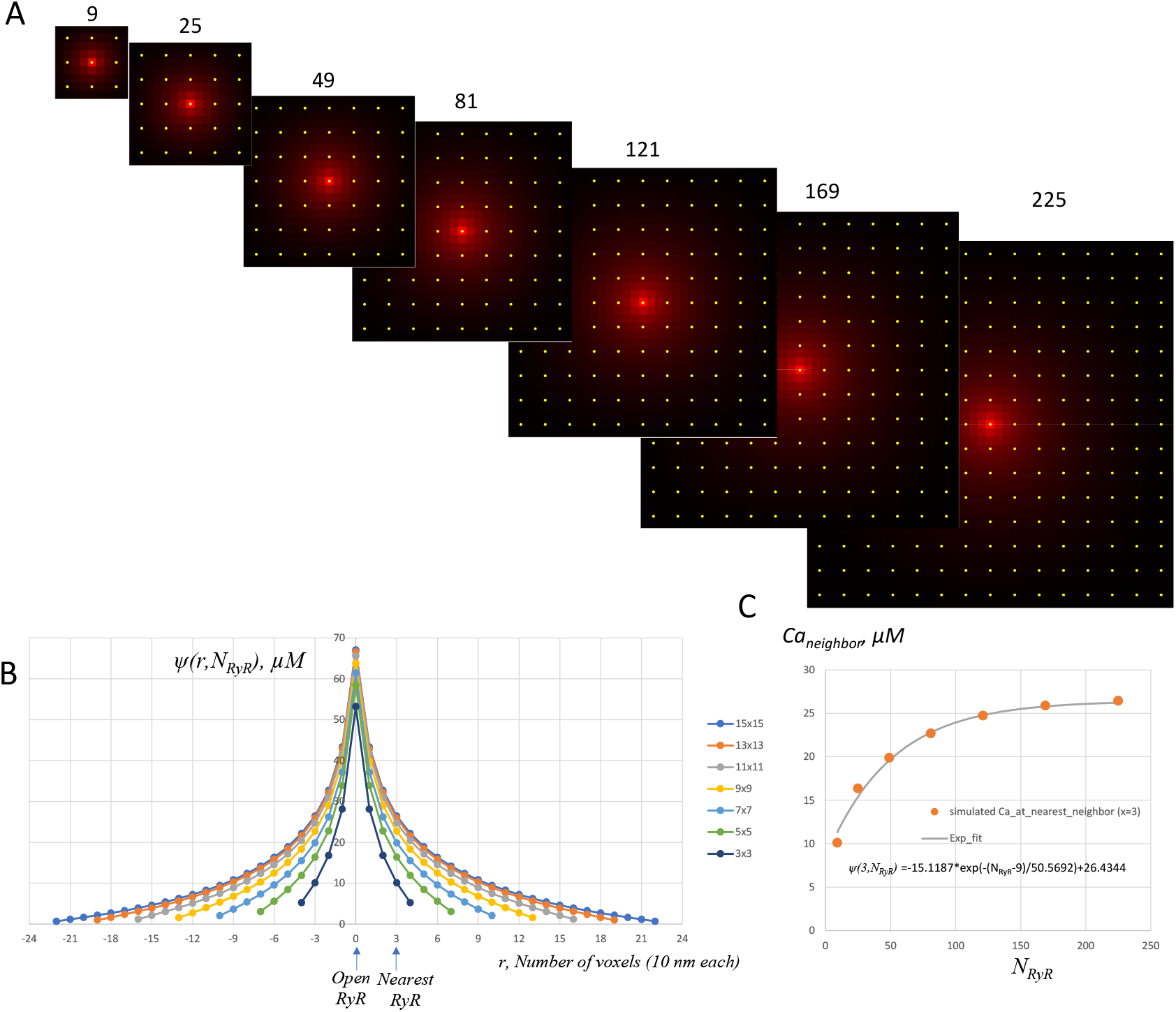
Effect of Ca^2+^ escape from dyadic space on RyR interaction profile in CRUs of different sizes (boundary effect). A: A cascade of distributions of Ca^2+^ in dyadic space for CRUs having different numbers of RyRs (shown by yellow circles): 3×3, 5×5, 7×7 etc. The distributions were simulated by running Stern at al. spark model [52] that describes function each individual RyRs within an individual CRU. The distributions were assessed 10 ms after one RyR channel opens in the middle of each CRUs. Other RyRs were not allowed to open. The initial [Ca] in JSR was set to 1 mM. [Ca] is coded by red shades, saturating (pure red) at 60 μM. The voxel size in the model is 10×10 nm in xy plane parallel to the cell surface membrane. B: Ca^2+^ profiles (interaction profiles) via the CRU center of the respective distributions in panel. The distance *r* is given in the model voxel size (10 nm). Arrows show positions of open RyR in the CRU center (r=0) and its nearest RyR neighbor (r=3 voxels). C. Plot of Ca^2+^ at the nearest neighbor for CRUs of different sizes Ca_neighbor_ = *ψ(3,N*_*RyR*_) together with its exponential fit (R^2^=0.976) shown by grey line with equation at the bottom of the plot.

